# Endogenous bystander killing mechanisms enhance the activity of novel FAP-specific CAR-T cells against glioblastoma

**DOI:** 10.1101/2023.02.21.529331

**Authors:** Wenbo Yu, Nga TH Truong, Ruhi Polara, Tessa Gargett, Melinda N Tea, Stuart M Pitson, Michaelia P Cockshell, Claudine S Bonder, Lisa M Ebert, Michael P Brown

## Abstract

**Objectives:** CAR-T cells are being investigated as a novel immunotherapy for glioblastoma, but clinical success has been limited. We recently described fibroblast activation protein (FAP) as an ideal target antigen for glioblastoma immunotherapy, with expression on both tumor cells and tumor blood vessels. However, CAR-T cells targeting FAP have never been investigated as a therapy for glioblastoma.

**Methods:** We generated a novel FAP targeting CAR with CD3ζ and CD28 signaling domains and tested the resulting CAR-T cells for their lytic activity and cytokine secretion function *in vitro* (using real-time impedance, flow cytometry, imaging, and bead-based cytokine assays), and *in vivo* (using a xenograft mimicking the natural heterogeneity of human glioblastoma).

**Results:** FAP-CAR-T cells exhibited target specificity against model cell lines and potent cytotoxicity against patient-derived glioma neural stem cells, even when only a subpopulation expressed FAP, indicating a bystander killing mechanism. Using co-culture assays, we confirmed FAP-CAR-T cells mediate bystander killing of antigen-negative tumor cells, but only after activation by FAP-positive target cells. This bystander killing was at least partially mediated by soluble factors and amplified by IL-2 which activated the non-transduced fraction of the CAR-T product. Finally, a low dose of intravenously administered FAP-CAR-T cells controlled, without overt toxicity, the growth of subcutaneous tumors created using a mixture of antigen-negative and antigen-positive glioblastoma cells.

**Conclusions:** Our findings advance FAP as a leading candidate for clinical CAR-T therapy of glioblastoma and highlight under-recognized antigen non-specific mechanisms that may contribute meaningfully to the antitumor activity of CAR-T cells.

## INTRODUCTION

Glioblastoma is the most lethal form of primary brain tumor and there is an urgent need for more effective therapies. Despite receiving intensive treatment upon initial diagnosis with all three classical pillars of cancer therapy (surgery, radiotherapy, and chemotherapy), patients almost inevitably relapse within a matter of months^1^. There is no standard treatment for recurrent glioblastoma, and most patients will die within 6 months of recurrence ^2^. The recent emergence of a ‘fourth pillar’ of cancer treatment, immunotherapy, may provide new hope that this dismal picture can be changed for patients with glioblastoma soon.

Chimeric antigen receptor (CAR)-T cells are one of the success stories of cancer immunotherapy, but so far clinical impacts are limited to B-cell malignancies. There remain significant hurdles to overcome before this success can be extended to solid tumors ^3^. One clear challenge is the lack of ideal target antigens in solid tumors. This sits in contrast to B-cell malignancies where lineage markers, such as CD19, show near-ubiquitous expression on cancerous cells and no expression outside the hematologic compartment. Nevertheless, the possibility of adapting CAR-T cell therapy to the treatment of glioblastoma is under extensive investigation ^4^. Most interest to date for clinical CAR-T targeting of glioblastoma has focused on targeting the antigens EGFRvIII, HER2 and IL-13R2α ^5-12^. Positive outcomes from clinical trials to date have generally been modest. However, some tumor regressions have been observed in recent studies, particularly using localized delivery of CAR-T cells, although such responses are generally transient ^7, 11, 12^. Given the profound tumor heterogeneity of glioblastoma, identification of more and better targets is a key focus of research. This should be coupled with the identification of innovative approaches to broaden CAR-T cell reactivity.

Fibroblast activation protein (FAP) is a surface-expressed proteolytic enzyme that is well known for its expression in cancer-associated fibroblasts ^13^. Because of this, FAP has been identified as a promising immunotherapy target for carcinomas which are typically heavily infiltrated with fibroblasts, such as those of prostate, breast, lung, and pancreas. In those studies, FAP has primarily been investigated as the target of tumor supporting stroma ^14-18^, but can also serve as a direct tumor target in certain cancers such as mesothelioma ^19, 20^. We and others found FAP to be consistently over-expressed in glioblastoma compared to normal brain, although we failed to find fibroblast expression of FAP in glioblastoma ^21, 22^. Rather, our detailed expression analyses of all cell types in glioblastoma revealed heterogeneous expression of FAP on the tumor cells, coupled with near-ubiquitous expression around tumor blood vessels, with expression observed on both endothelial cells and pericytes. These studies suggest FAP is an ideal immunotherapy target for glioblastoma, as it should allow targeting of not only tumor cells, but also the tumor’s supporting vascular networks ^21^.

Several FAP-specific CARs have been described in the literature. All FAP-CARs have been derived from one of four mouse monoclonal antibodies (mAbs). A CAR developed from the FAP-5 mAb cross-reacts with both mouse and human FAP. However, FAP-5-CAR-T cells showed serious toxicities in mice and this CAR has not been pursued further ^23^. A second CAR developed from the F19 mAb recognizes only human FAP, thus this CAR could not be used to test potential host FAP-related toxicities in mouse models ^19, 20, 24^. A third CAR developed from mAb 73.3 cross-reacts with both mouse and human FAP and showed limited toxicities in mice ^16, 17^. CAR-T cells derived from both F19 and 73.3 mAbs showed antitumor effects in mouse models of multiple types of transplanted tumors ^16, 20^ and have now progressed into phase 1 clinical trials (ClinicalTrials.gov Identifiers: NCT039325652019 and NCT017221492015). However, neither CAR-T has been tested on glioblastoma. F19-CAR-T cells were well tolerated clinically ^25^, although no detailed report of either trial has been released yet.

In this study, we generated a FAP-specific CAR derived from the MO36 mAb, which was selected from phage display and recognizes both murine and human FAP ^26^. Another CAR based on MO36 previously showed antitumor effects in a lung cancer mouse model without toxicity ^15^. We tested FAP-CAR-T cell function using model cell lines, tumor cells expanded from patient glioblastoma tissues, and a mouse model designed to recapitulate the natural heterogeneity of glioblastoma. In addition to confirming the specificity and activity of FAP-CAR-T cells, we also uncovered a potent bystander killing mechanism that could facilitate more complete tumor destruction than would be predicted by antigen expression alone.

## RESULTS

### Generation and production of CAR T cells targeting FAP

Based on previous studies ^15, 26^, we designed and constructed a FAP-CAR comprising the MO36 single chain fragment-variable (scFv), a myc epitope tag, a long spacer, and CD28 and CD3ζ signaling domains (Figure. 1a). This CAR transgene was integrated in CD3/CD28-activated human T cells via lentiviral transduction. CAR expression on transduced T cells was assessed by flow cytometry (Figure. 1b). Transduction efficiency values sufficient to test CAR-T cell function (mean 25.1%, SD 7.2% from transduction of 8 donors) were achieved using unconcentrated viral supernatant.

**Figure 1.**
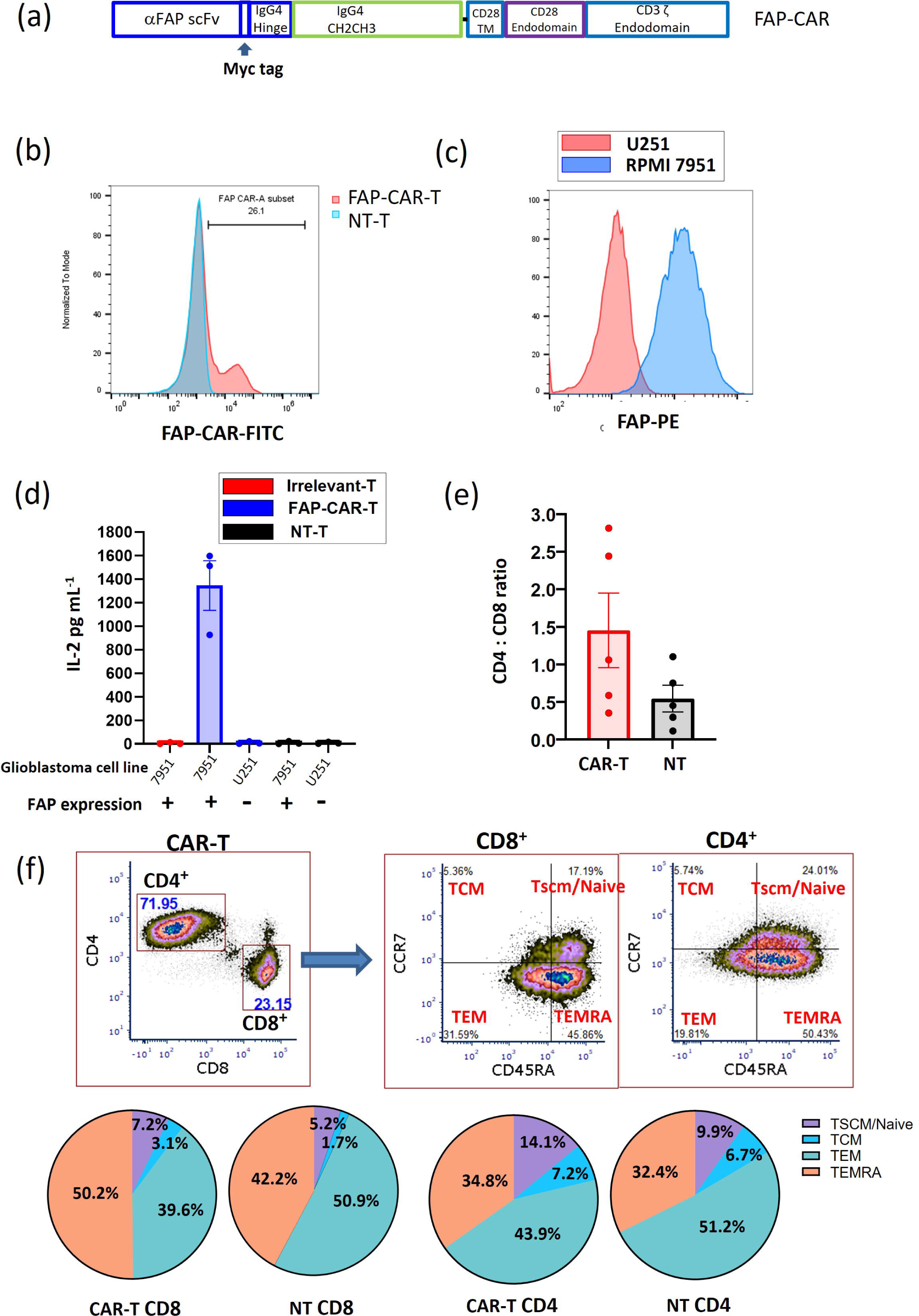
Structure and characterization of the FAP-CAR. **(a)** Schematic representation of the FAP CAR construct. The CAR consisted of the MO36 anti-FAP scFv linked to a CD28 transmembrane (TM) and CD28 and CD3ζ costimulatory endodomains. The FAP-CAR contains the myc tag at the end of the scFv and a spacer from human IgG4 CH2CH3 fragment with several point mutations. **(b)** CAR expression on CAR-T cells was detected by flow cytometry. The CAR-T cells were stained by FITC-conjugated anti-myc tag antibody. One representative sample from 8 donors is shown. **(c)** FAP expression on target cell lines was confirmed by flow cytometry. **(d)** CAR-transduced or non-transduced T cells were co-cultured with target cells with or without FAP expression and the IL-2 concentration in supernatants determined 48h later. The data represent the mean ± SE from three independent assays. Each assay represents the average value from duplicate wells. Anti-EphA3 CAR-T cells provided the irrelevant CAR in this assay. (e-f): FAP-CAR-T cell products were manufactured from 5 different healthy donors and analysed by flow cytometry, using an antibody to the myc tag to identify the CAR^+^ (transduced) cells. Shown in **(e)** is a summary of the CD4:CD8 ratio within the CAR^+^ and CAR^-^ populations, while **(f)** depicts the memory phenotype; pie charts show mean values across the 5 donors. T cells were categorized as TSCM/Naïve (CCR7^+^/CD45RA^+^), TCM (CCR7^+^/CD45RA^-^), TEM (CCR7^-^/CD45RA^-^), and TEMRA (CCR7^-^/CD45RA^+^).

CAR-T cell responses to FAP stimulation *in vitro* were initially measured by co-culturing CAR-T cells with FAP-expressing long-term cancer cell lines. The RPMI-7951 melanoma cell line, which expresses uniformly high levels of FAP, was used for initial testing of FAP-CAR-T cells, while the FAP-negative U251 glioblastoma cell line was used as a negative control (Figure 1c). These tumor cell lines were co-cultured with FAP-CAR-T cells or CAR-T cells targeting an irrelevant antigen (EphA3) for 48 hours, and the level of IL-2 in the culture supernatant was measured by ELISA (Figure 1d). The data indicate that FAP-CAR-T cells (but not irrelevant CAR-T cells) responded specifically to target cells expressing FAP, whereas there was no response to cells lacking FAP. Further analysis of the CAR-T cells from 5 donors revealed a relatively balanced CD4:CD8 ratio and dominant populations of effector memory (TEM) and terminally differentiated effector (TEMRA) cells within the cell products (Figure 1 e-f).

### Cytotoxicity of FAP-CAR-T cells against glioblastoma cell lines and patient-derived glioma neural stem (GNS) cells

We next used impedance-based real-time analysis to assess FAP-CAR-T cell cytotoxicity. As target cells, the FAP-expressing U87 human glioblastoma cell line was compared with the FAP-negative glioblastoma cell line U251 (Figure 2a). As expected, FAP-CAR-T cells were strongly cytotoxic against U87 target cells, whereas non-transduced T cells (NT-T) induced only minimal background lysis. FAP-CAR-T cells failed to lyse U251 cells above background levels, thereby confirming the specificity of the CAR.

**Figure 2.**
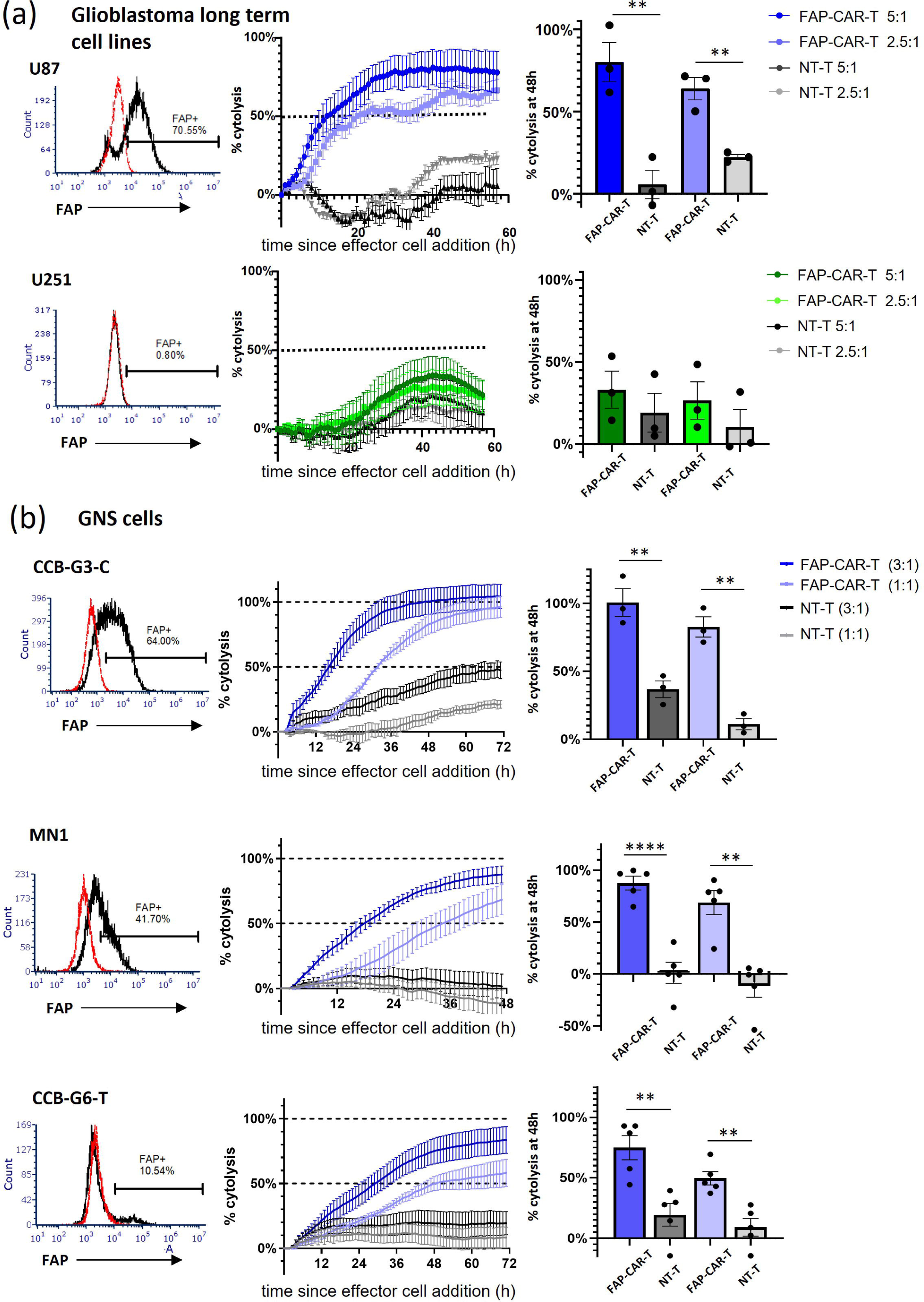
FAP-CAR-T cells display cytotoxic activity against glioblastoma cell lines and GNS cells expressing varying levels of FAP. Left panels: FAP expression on target cells was examined by flow cytometry. Red dotted lines indicate isotype control and black solid lines indicate anti-FAP antibody. Centre and right panels: Target cells were established in a CytoView-Z plate and placed in the Maestro Z system for up to 24h, followed by addition of FAP-CAR-T cells as effectors or non-transduced T cells (NT-T) as a control. Cytotoxicity was monitored over time and represented as percentage of target lysis averaged across duplicate wells. Representative examples shown in centre panels and % cytolysis at 48h after effector cell addition is plotted in the right panels. The effector:target (E:T) ratio is indicated. *t*-test; * *P* < 0.05, ** *P* < 0.01. (a) FAP^+^ U87 and FAP^-^ U251 cells were used as target cells. The data represents the mean ± SE from three independent assays. Each assay represents the average value from duplicate wells. (b) GNS cells expressing varying levels of FAP were used as target cells. The data represent the mean ± SE from 3-5 independent assays. Each assay represents the average value from duplicate wells.

Although long-term established cell lines such as U87 provide a suitable model to test a diverse range of therapeutic approaches, these cells may not adequately represent primary glioblastoma cells. Therefore, we next selected a panel of patient-derived glioma neural stem (GNS) cells as a more biologically relevant type of target cell to test the cytotoxic activity of our FAP-CAR-T cells. We checked a panel of different GNS cells for FAP expression and chose three cultures which express different levels of FAP. CCB-G3-C, MN1, and CCB-G6-T are representative of high, medium, and low level FAP expression, respectively (Figure 2b). We found substantial killing of all of three GNS cell lines (Figure 2b). Of note, when cultured with MN1 cells, which are <50% FAP^+^, the FAP-CAR-T cells induced near-complete cytotoxicity (> 90% lysis). Even more remarkably, in the case of CCB-G6-T cells, even a 10% FAP^+^ population was sufficient to induce the lysis of >80% of all target cells.

These observations suggest that FAP-CAR-T cells are capable of potent bystander killing, whereby activation of CAR-T cells by a subpopulation of antigen-expressing target cells leads to killing of antigen-negative tumor cells as well ^27^. This idea was further supported by a simple flow cytometry assay, wherein CCB-G6-T target cells were co-cultured with FAP-CAR-T cells for 48 hours and the residual viable cells assessed for FAP expression (Supplementary Figure 1). After this time, the small FAP^+^ population within CCB-G6-T was significantly depleted, as expected. Notably, however, the loss in cell viability amongst the entire target cell population was ∼50%, far more than the FAP^+^ population frequency, further supporting a bystander killing effect of FAP-CAR-T cells. The observed loss of FAP expression was unlikely to be due to trogocytosis (27), as we did not detect any transfer of FAP to CAR-T cells (data not shown). However, this assay did not allow definitive tracking of the fate of FAP^+^ versus FAP^-^ target cells, and so we sought a better system to address this issue more comprehensively.

### Confirmation that FAP-CAR-T cells induce antigen-dependent bystander killing of tumor cells

To confirm the bystander killing capability of FAP-CAR-T cells, we labeled FAP^+^ GNS cell lines with a fluorescent dye (CellTrace Far Red; CTFR) and mixed them with unlabeled GNS cells lacking FAP expression. This target-cell mixture was incubated with FAP-CAR-T cells or NT-T cells for 48 hours and the viability of each target-cell population analyzed by flow cytometry (see Supplementary Figure 2 for an overview of the experimental design). Two different pairs of FAP^+^/FAP^-^ GNS cell lines were used. Pair 1 consisted of FAP^+^ MN1 cells with FAP^-^ RKI1 cells, and Pair 2 consisted of FAP^+^ CCB-G3-C cells with FAP^-^ SJH1 cells. Whereas MN1 and CCB-G3-C cells showed uniform expression of FAP by flow cytometry (see Figure 2b), RKI1 and SJH1 cells lacked any detectable surface expression (Figure 3a). These FAP-negative GNS cells also lacked significant gene expression by microarray analysis (data not shown).

**Figure 3:**
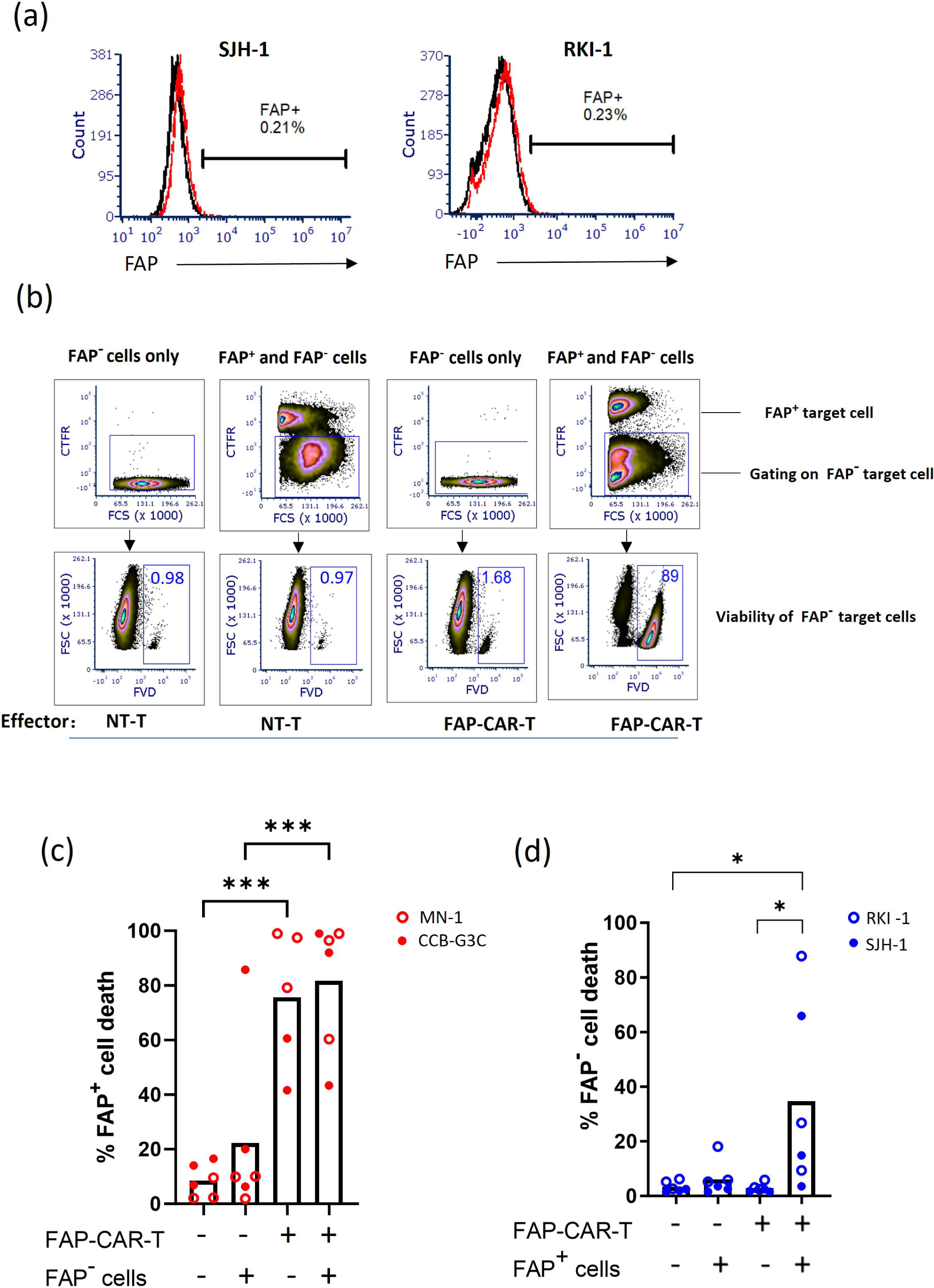
FAP-CAR-T cells exert a bystander killing effect on FAP-tumor cells after CAR engagement. FAP ^+^ tumor cells were stained with cell trace far red (CTFR) for discrimination and then FAP^+^ and FAP^-^ tumor cells were mixed at a 1:1 ratio, followed by treatment with either FAP-CAR-T cells or non-transduced T cells (NT-T) at a 1:1 E:T ratio. Single cultured target cells were treated in the same way as control. The viability of target cells was analyzed 48h later by flow cytometry using fixable viability dye (FVD). **(a)** FAP expression on target cells was examined by flow cytometry. The FAP^-^ SJH-1 and RKI-1 GNS cell lines are shown here; refer to Fig 2 for FAP ^+^ MN1 and CCB-G3-C cells. Red dotted lines indicate isotype control and black solid lines indicate anti-FAP antibody. **(b)** Gating strategy of a representative experiment (Pair 1) showing cell death within the FAP^-^ target cell population. Refer to Supplementary Figure 3 for representative examples of cell death within the FAP^+^ target cell population. **(c-d)** The percentage of cell death of FAP^-^ GNS cells (**c**) or FAP^+^ GNS cells (**d**) after 48h co-culture was calculated. Data were pooled from three independent experiments, including both Pair 1 and Pair 2 samples (total n = 5-6). Samples from both Pair one and Pair two were pooled together. *t*-test; * *P* < 0.05, *** *P* < 0.001.

An example of the gating strategy and analysis of the FAP^-^ target cell fraction is shown in Figure 3b, while pooled data are summarized in Figure 3(c-d). As expected, FAP-CAR-T cells induced efficient killing of FAP^+^ target cells (Fig 3c, and representative data in Supplementary Figure 3). Also as expected, the viability of FAP^-^ tumor cells cultured alone was not affected by incubation with FAP-CAR-T cells, thereby ruling out off-target effects. But when FAP^-^ target cells were mixed with FAP^+^ target cells and incubated with FAP-CAR-T cells, significant killing of the FAP^-^ tumor cells was observed (Figure 3b and d). Thus, FAP-CAR-T cells can exert substantial bystander killing of tumor cells lacking FAP, but only once they have been activated by cells expressing their cognate antigen. To determine if the bystander killing effect is unique to FAP-CAR-T cells, we also tested GD2-specific CAR-T cells on two GNS cells expressing low levels of GD2 (16-26% GD2-positive). Real-time cytotoxicity assays revealed complete lysis of these target cells despite heterogeneous antigen expression (Supplementary Figure 4), thus suggesting that the bystander killing activity of CAR-T cells is not restricted to a particular CAR.

### Bystander cytotoxicity occurs independently of Fas/FasL but can be mediated by soluble factors

Bystander killing of antigen-negative targets has recently been reported to occur via Fas-FasL interactions^27^. To determine if this molecular axis was at play in our system, we first investigated expression of Fas on our GNS cell lines, and expression of FasL on our FAP-CAR-T cells. Although variable levels of Fas were detected on the GNS cell lines (RKI1, SJH1 and CCB-G6-T) found to be susceptible to bystander killing (Supplementary Figure 5a), there was almost no expression of FasL by FAP-CAR-T cells, even after co-culture with FAP^+^ target cells and with permeabilization to detect intracellular stores of FasL (Supplementary Figure 5b). Soluble FasL in culture medium was also low (< 20 pg mL^-1^, data not shown). ZB4 was reported as a function-blocking anti-Fas antibody^28^. We confirmed this in our soluble FasL induced cytotoxicity assay (Supplementary Figure 5c). However, ZB4 failed to attenuate either direct killing (data not shown) or bystander killing (Supplementary Figure 5d-e). We therefore conclude that bystander killing is unlikely to be mediated by Fas/FasL interactions in our system.

We next hypothesized that soluble factors such as cytokines could contribute to the bystander killing effect. This was tested by separating antigen-negative target cells from CAR-T cells using a semi-permeable membrane. FAP-CAR-T or NT-T cells were seeded together with FAP^+^ GNS cells in the lower chambers of Transwell assemblies, while FAP^-^ GNS cells were seeded in the upper chambers (Figure 4a). After 48 hours, cells in the upper chambers were stained using propidium iodide (PI) to label dead cells and Hoechst 33342 to visualize total nuclei, and cell viability calculated by the ratio of PI/Hoechst-stained cells. The same combinations of FAP^-^ SJH-1/FAP^+^ CCB-G3-C cells and FAP^-^ RKI-1/FAP^+^ MN-1 cells from Figure 3 were used in this assay. In keeping with our hypothesis, significant killing of FAP^-^ tumor cells in the upper chamber (SJH-1 or RKI-1) was observed when FAP-CAR-T cells were present in the lower chamber together with FAP^+^ tumor cells (Figure 4b-d).

**Figure 4:**
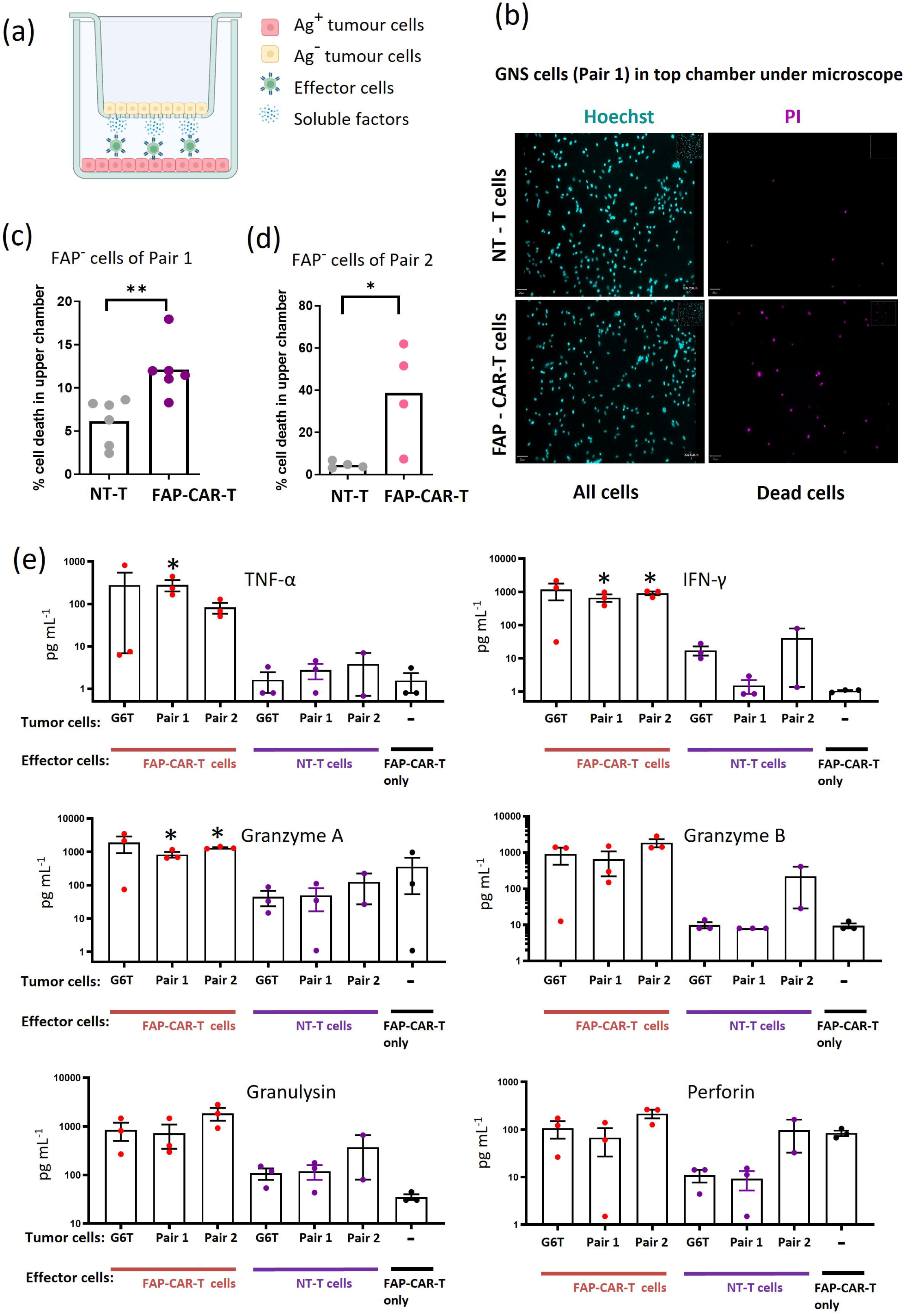
Bystander killing by FAP-CAR-T cells can be mediated by soluble factors. **(a)** Schematic of assay set-up. **(b)** Representative fluorescence images of Hoechst 33258 (left) and PI (right) staining on FAP^-^ tumor cells cultured in upper chambers for 48h. Effector cells (NT-T cells or FAP-CAR-T cells) added to lower chambers are indicated. Images were collected using a 10X objective. **(c-d)** Quantification of percent of FAP^-^ target cell death in upper chamber. Shown in c is FAP^-^ RKI-1 from Pair 1, n = 6, pooled from 3 experiments. Shown in **d** is FAP^-^ SJH-1 from Pair 2, n = 4 pooled from 2 experiments. Each dot represents the mean percentage of cell death from 3 fields of view of each well. *t*-test; * *P* < 0.05, ** *P* < 0.01. **(e)** Concentrations of secreted factors in supernatants of cocultures were determined by Legendplex kit. FAP-CAR-T stimulated by the FAP^+^ GNS cell line CCB-G6-T was used as positive control. Supernatants were collected after 48h. Data are pooled from 3 experiments except NT-T of Pair 2 which only have two data points. Mann-Whitney *U*-tests; * *P* < 0.05.

Since the Transwell experiments suggested that bystander killing of antigen-negative cells can be at least partly mediated by soluble factors, a panel of cytokines and cytotoxic proteins was measured in co-culture supernatants using a multiplexed flow cytometric bead assay. Large amounts of TNF-α, IFN-γ, granzyme A, granzyme B and granulysin were released when FAP-CAR-T cells were co-cultured with FAP^+^ GNS cells, but not when cultured alone (Figure 4e). Perforin was also detected in the co-culture supernatants but was similarly secreted in the absence of target cells, suggesting constitutive secretion by FAP-CAR-T cells. As expected, the NT-T cells generally secreted only low levels of these factors, even in the presence of FAP^+^ GNS tumor cells. Together, our results indicate that FAP-CAR-T cells can be activated by antigen to secrete a range of cytotoxic cytokines and proteins, which may contribute to the contact-independent bystander killing observed in the Transwell system.

### FAP-CAR-T cell activation mobilizes the CAR-negative T-cell fraction to enhance cytotoxicity

It has long been recognized that, in addition to antigen-dependent cytotoxicity, T cells can also undergo cytokine-induced bystander activation ^29^. Our FAP-CAR-T cell products contain ∼75% CAR-negative T cells, leading us to hypothesize that CAR-T cells may secrete cytokines to mobilize the non-transduced T cells into the antitumor response after activation by FAP-expressing tumor cells (Figure 5a). In this way, the whole population of T cells could be synchronized to exert cytotoxicity against both antigen-positive and antigen-negative tumor cells. Cytokines including IL-2, IL-12, IL-15 and IL-21 have been reported to initiate antigen nonspecific T-cell activation^30-32^. And although IL-15 and IL-12 are primarily produced by antigen presenting cells, IL-2 and IL-21 can be produced by T cells. We showed that IL-2 was secreted by FAP-CAR-T cells after antigen activation (Figure 1d), suggesting that it could be one of the cytokines to mobilize normal T cells during bystander killing.

**Figure 5.**
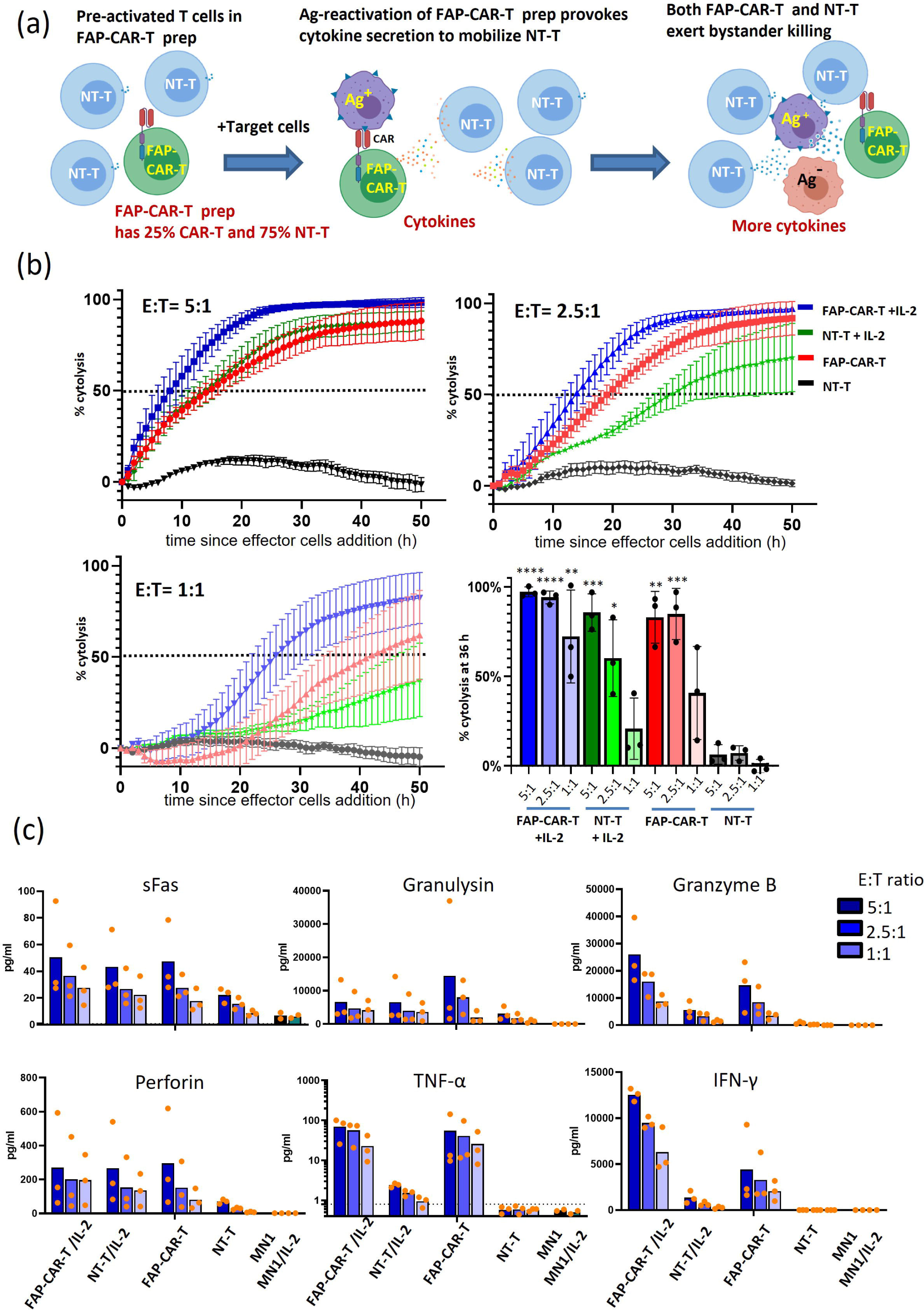
IL-2 induces T cells to exert antigen nonspecific cytotoxicity against GNS cells. **(a)** A conceptual diagram of the theory. After antigen-specific activation, FAP-CAR-T cells secrete IL-2 which mobilizes NT-T cells to exert further cytotoxicity against tumor cells. **(b)** Cytotoxicity assay of FAP-CAR-T or NT-T cells against MN1 GNS cells, with or without exogenous IL-2 (20ng/ml). FAP^+^ MN1 cells were established in a CytoView-Z plate for 24h before addition of effector cells. Target cell growth was monitored by impedance values in real time by the Maestro Z system. Time-course graphs for varying E:T ratios, as indicated, show target lysis as % cytotoxicity. A summary of % cytolysis at 36h after effector cell addition is shown bottom right. Data points represent mean ± SEM from 3 independent experiments. Each assay represents the average value from duplicate wells. **(c)** Secreted factors were measured in supernatants 50h after addition of effector cells using Legendplex assay. A total of 12 analytes were measured in 3 experiments (or 2 experiments for MN-1 and MN-1/IL-2 controls). Results of 6 key cytotoxic factors are shown, with the other 6 being depicted in Supplementary Figure 6. Statistical analysis of these data is presented in Table 1.

**Table 1.** Linear fitting regression was performed using cytolysis %_36h_ vs cytokine concentration _(pg mL_^-1^_)_. The cytokines were ranked by their adjusted R^2^ value from high to low in the table. *P-*value was calculated by ANOVA. The significance of differences between NT-T ± IL-2 or between NT-T and FAP-CAR-T was determined by paired *t*-test. *P* < 0.05 was considered significant.

| Cytokine | Correlation with cytotoxicity |  | IL-2/N-T vs N-T | IL-2/N-T vs IL-2/FAP-CAR-T |
| --- | --- | --- | --- | --- |
|  | Adj-R2 | P-value | (Significant or not) | (Significant or not) |
| Granzyme B | 0.47994 | 8.66E-07 | Yes | Yes |
| IL-4 | 0.43468 | 4.06E-06 | Yes | Yes |
| IFN $\gamma$ | 0.43145 | 4.51E-06 | Yes | Yes |
| Perforin | 0.29523 | 2.52E-04 | Yes | **No** |
| IL-6 | 0.26234 | 5.99E-04 | Yes | Yes |
| TNFα | 0.25307 | 7.60E-04 | Yes | Yes |
| IL-10 | 0.22098 | 0.00171 | **No** | Yes |
| IL-17a | 0.20964 | 0.00226 | **No** | Yes |
| Granulysin | 0.14905 | 0.00962 | Yes | Yes |
| sFas | -0.02626 | **0.81865** | Yes | Yes |
| sFas L | -0.02713 | **0.88065** | Yes | Yes |

We performed additional real-time cytotoxicity assays to explore these concepts by recapitulating the bystander activation and killing processes *in vitro*. The cytotoxicity of FAP-CAR-T cells or NT-T cells was tested against MN1 GNS cells expressing moderate levels of FAP, with or without the addition of 20ng/ml exogenous IL-2 (Figure 5b). FAP-CAR-T cells displayed strong cytotoxicity against MN1 as expected, which was further boosted by the addition of IL-2. This effect was marginal at high E:T ratios but became more pronounced at lower E:T ratios as CAR-expressing effector cells became limiting. Of note, IL-2 also dramatically increased the cytotoxicity of control T cells. For example, at a high (5:1) E:T ratio, NT-T cells supplemented with IL-2 induced as much tumor cell killing as FAP-CAR-T cells. This observation supports the concept that, although IL-2 may also boost the activity of antigen-activated CAR^+^ cells, it has the greatest impact on the non-transduced ‘bystander’ cells.

We further analyzed cytokines and cytotoxic proteins in supernatant samples collected from these assays. In addition to IL-2, we measured another 11 analytes (Figure 5c, Supplementary Figure 6, Table 1). The addition of IL-2 to NT-T cells significantly increased the levels of all factors except IL-10 and IL-17a (see column 3 in Table 1), suggesting that these two cytokines were upregulated only by CAR stimulation, not IL-2. Furthermore, by comparing NT-T cells + IL-2 to CAR-T cells + IL-2, we could show that all factors were further boosted by CAR engagement except for perforin (see column 4 in Table 1), suggesting that IL-2 alone was sufficient to induce maximum secretion of soluble perforin. Interestingly, all analytes showed a positive correlation with the extent of cytotoxicity except soluble Fas and FasL (Table 1, column 2). These results agree with those from our ZB4 blocking experiment (Supplementary Figure 5), which did not support a significant role for the Fas/FasL axis in cytotoxicity in our system.

In summary, our findings suggest that cytokines, such as IL-2 used here, can stimulate the non-transduced fraction of a CAR-T cell product also to mediate tumor cell killing. Furthermore, this amplification of the response may be an important aspect of the bystander killing mechanism, because IL-2 also resulted in upregulated secretion of factors such as TNFα, IFNγ, granzyme and granulysin, which are candidate mediators of bystander killing. Other cytokines upregulated in this system (IL-4, IL-6, IL-10, and IL-17) are not directly involved in cytotoxicity but can support T- and B-cell function.

### FAP-CAR-T cells exhibit a lack of toxicity against healthy cells and control the growth of tumors in a subcutaneous glioblastoma xenograft model despite heterogeneous antigen expression

To assess the function of our CAR-T cells *in vivo*, we tested their ability to control the growth of subcutaneous human xenograft tumors in immunodeficient (NSG) mice. Glioblastoma is well-known for its cellular heterogeneity, and we have shown that expression of FAP by glioblastoma tumor cells is also heterogenous ^21^. Accordingly, we generated a model with mixed FAP^+^ and FAP^-^ tumor cells. FAP^+^ U87 cells were engineered to express RFP, while FAP^-^ U251 cells were engineered to express GFP, to allow monitoring of the mixed tumor cell populations. In an initial *in vitro* assay, the single target cell populations or the target cell mixture were co-cultured with FAP-CAR-T cells or NT-T cells and surviving cells were counted using fluorescence microscopy 24 hours later. Counts of FAP^-^ U251 cells were reduced significantly only when co-cultured with both U87 cells and FAP-CAR-T cells (Figure 6a), suggesting that antigen-negative U251 tumor cells are also susceptible to bystander killing from FAP-CAR-T cells, similar to our earlier observations using GNS cells. Thus, in addition to replicating the heterogeneous nature of glioblastoma tumors, a mixed U87/U251 tumor model also seems suitable to examine the bystander killing capacity of FAP-CAR-T cells *in vivo*.

**Figure 6.**
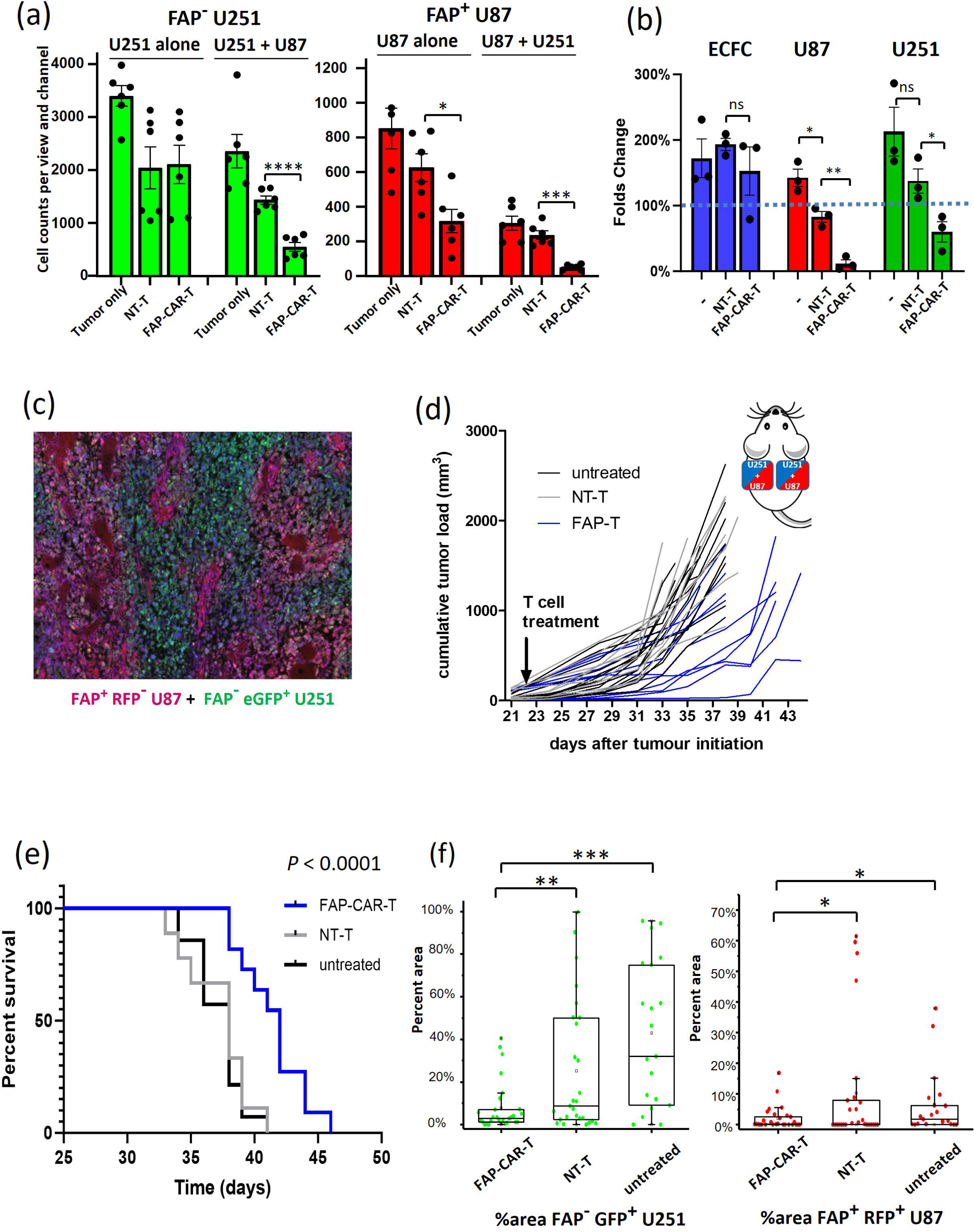
FAP-CAR-T cells control tumor growth in a mouse model with heterogeneous antigen expression. **(a)** *In vitro* bystander killing assay of FAP^-^ U251-GFP and FAP^+^ U87-RFP. Cells were cultured individually or together and were treated with FAP-CAR-T or NT-T cells. Fluorescence images were taken 24h later and GFP and RFP positive cells were counted using ImageJ. Graphs show mean and SE from 6 counts (3 fields of views × 2 repeats). Effector cells derived from one donor were used **(b)** ECFCs were co-cultured with an equal number of U87-RFP and U251-GFP cells and subsequently treated with effector cells (either FAP-CAR-T or NT-T cells) or left untreated (indicated as -). After 48 hours, cell counts were determined using flow cytometry and fold change from baseline calculated. The assay was performed three times, with FAP-CAR-T or NT-T cells derived from three different donors. Graphs show mean and SEM. **(c-f)** Subcutaneous tumours were created on both flanks of NSG mice using a mixture of FAP^-^ U251-eGFP and FAP^+^ U87-RFP cells (as illustrated in the inset to **d**). Tumors were allowed to establish for 21 days and then mice were treated with a single intravenous injection of 1 x 10^6^ FAP-CAR-T cells (blue) or NT-T cells (grey) or left untreated (black). FAP-CAR-T group n = 11, NT-T group n = 9, untreated group n = 14. (**c**) Representative tumor tissue section showing the balance of U251 (green) and U87 (red) cells, with DAPI staining to visualize nuclei. **(d)** Tumor growth curves for each mouse. The volume of tumors on each flank was measured every 2-3 days and summated to calculate total tumor load per mouse. Tumor growth in FAP-CAR-T group was significantly lower than untreated or NT-T groups (both adjusted *P* < 0.0001; Type II ANOVA using TumGrowth analysis). **(e)** Animals were humanely killed once at least one tumor reached 1000 mm^3^ and survival analysis performed using a Kaplan–Meier plot. *P* < 0.0001 by Log-Rank test. **(f)** GFP and RFP area of endpoint tumor sections were calculated. Data are pooled from a total of 28 sections from 13 tumors of FAP-CAR-T treated group, 27 sections from 13 tumors of NT-T treated group and 20 sections from 9 tumors of untreated group. “▢” represents mean; “x” represents minimal and maximal data; Bar represent 1% and 99% data distribution; Box represents 25%, 50% and 75% data distribution. *t*-test; * *P* < 0.05, ** *P* < 0.01, *** *P* < 0.001

Another important consideration before proceeding to animal studies was safety. We needed to determine whether bystander killing would result in off-target toxicity by targeting normal cells. To address this concern, we conducted a four-way co-culture experiment. Primary human Endothelial Colony Forming Cells (ECFCs), derived from circulating progenitor cells in peripheral blood, serve as a model for healthy blood vessels^33^, and as shown in Supplementary Figure 7, express minimal FAP protein in their surface. We co-cultured ECFCs with U87 and U251 cells and subsequently treated the cell mixture with effector cells (either FAP-CAR-T or NT-T cells), then 48 hours later flow cytometry was used to assess relative numbers of the various target cell populations (Figure 6b). As expected, FAP-CAR-T cells killed the FAP^+^ U87 cells. The FAP^-^ U251 cells were also killed, in keeping with bystander killing activity. However, ECFCs in the same culture well were not targeted, suggesting that the key bystander killing mechanisms at play in this system only target malignant cells.

The fluorescently tagged U87 and U251 cell lines were mixed at a 2:1 ratio, respectively, before subcutaneous implantation in each flank of NSG mice. The 2:1 mixing ratio was selected after an initial pilot study and resulted in the most evenly balanced tumors at endpoint (Figure 6c, Supplementary Figure 8 a-b). Compared to either NT-T cells or no treatment, intravenous administration of a low dose (1 x 10^6^) of FAP-CAR-T cells significantly reduced tumor growth (adjusted *P* < 0.0001 by Type II ANOVA for both control comparisons; Figure 6d), and significantly increased survival time of treated mice (both *P* < 0.0001 by Log-Rank test; Figure 6e). Notably, the FAP-CAR-T cells did not induce any overt signs of toxicity, with no mice needing to be culled due to weight loss, signs of distress/illness, or poor body condition. In mice treated with FAP-CAR-T cells, both the GFP and RFP signals in necropsied tumor tissues were significantly reduced compared to tumors in either control group (Figure 6f), suggesting CAR-T cell targeting of both FAP^+^ U87-RFP and FAP^-^ U251-GFP cells. These findings support the effectiveness of our FAP-CAR-T cells in controlling the growth of glioblastoma cells *in vivo*, even in the face of heterogeneous expression of target antigen.

## DISCUSSION

For many years, FAP has been recognized as a promising immunotherapy target for carcinomas and mesothelioma ^14-17, 19, 20^. More recently, we have proposed FAP as a candidate immunotherapy target antigen for glioblastoma ^21^. Studies from our group and others have shown that, in glioblastoma, FAP is heterogeneously expressed on tumor cells themselves, but more consistently by tumor-supporting stromal populations including endothelial cells and pericytes. This sits in sharp contrast to the limited expression in healthy tissues and vessels ^21, 34^. Despite all these characteristics, until now FAP has never been tested as a target antigen for immunotherapeutic strategies in glioblastoma.

Glioblastoma has striking intra-tumor heterogeneity ^35^. For initial confirmation of targeting specificity, we used long-term cell lines including U87 and U251, and these studies successfully demonstrated that our FAP-CAR-T cells secrete IL-2 and induce cytolysis only when FAP is expressed. However, these cell lines (which have been cultured for many decades *in vitro* in the presence of xenogeneic serum) may lose multipotency, develop pro-survival mechanisms and fail to model the disease properly ^36^. For further *in vitro* testing, we therefore moved to the use of glioma neural stem (GNS) cell lines, which are patient glioblastoma tissue-derived cells expanded under serum-free conditions to a limited passage number (generally < 30) and maintain a more *in vivo*-like phenotype ^21, 37^. Our group has developed a panel of GNS cells from patient samples and characterized them for FAP expression ^21^ and we used these GNS cells, together with those from the Q-Cell resource ^38^ as *in vitro* models to investigate more authentically how our CAR-T cells tackle the heterogeneity of glioblastoma, including low and heterogeneous patterns of FAP expression.

Our results using GNS cells in a sensitive real-time cytotoxicity assay indicated that the percentage of cells lysed by FAP-CAR-T cells can greatly exceed the percentage of target cells expressing the antigen. Further flow cytometry-based analysis of co-cultured GNS cells confirmed that FAP-CAR-T cells can kill antigen-negative tumor cells, but only in the presence of antigen-positive tumor cells. Antigen/epitope spreading is a phenomenon whereby immune reactivity is broadened following the release of antigens during chronic inflammation, and is dependent on an intact immune system and the involvement of antigen-presenting cells (APCs)^39^. It is highly unlikely that this phenomenon occurs in the reductionist co-culture systems used here. Rather, we conclude that FAP-CAR-T cells are capable of ‘bystander’ killing, a phenomenon long recognized in T-cell immunology but poorly understood ^40, 41^. The significance of bystander killing in CAR-T cell function is also just beginning to be appreciated ^27, 42^, and many questions remain. For example, it is unclear whether all CAR-T cell products have bystander killing capacity, or does this depend on factors such as CAR structure or target cell susceptibility? Based on our observations using CAR T cells targeting FAP or GD2, we believe that this function may be a common feature of CAR-T function that has been generally underappreciated in the field because it will not be revealed in such short-term cytotoxicity assays as the classical chromium release assay. By using the novel Maestro Z instrument to track target cell destruction over several days, we allow CAR-T cells the time required to be activated by CAR engagement and then deploy a range of cytotoxic mechanisms against antigen-negative tumor cells. Therefore, it is possible that all CAR-T cell products can perform bystander killing to some degree, but a sufficient observation period is required to record this phenomenon.

Another important question regarding bystander killing is: what are the key effector molecules? The major killing mechanism described for CAR-T cells has been direct cytolysis, which is induced by perforin and granzyme and delivered via a CAR-mediated immunologic synapse ^43^. Recent studies have suggested that the Fas/FasL signaling axis participates in CAR-T cell cytotoxicity, and also in bystander killing ^27, 42^. In contrast, we did not find evidence that Fas/FasL contributes to bystander killing (or cytotoxicity in general) by FAP-CAR T cells. We did, however, observe that the bystander killing effect was at least partly mediated by soluble factors, as significant cytotoxicity was observed even when the CAR-T cells and tumor cells were not in direct contact. Determining exactly which secreted factor or factors drive bystander killing by FAP-CAR-T cells is beyond the scope of the present study. However, we did detect secretion of a range of cytokines and cytotoxic granule components upon antigen engagement of FAP-CAR-T cells. Two cytokines of particular interest are TNFα and IFNγ, both of which have a direct antitumor effect ^44, 45^. Moreover, TNF has previously been shown to play a critical role in the antitumor activity of HER2-targeting CAR-T cells, and to facilitate bystander killing in the context of patient-derived colorectal tumoroids^46^. In this system, the relatively low penetrance of HER2-CAR-T cells meant that effective cytotoxicity could only be achieved with a diffusible factor such as TNF. We also detected secretion of surprisingly high levels of molecules such as granzyme B and perforin, which are normally involved in cytotoxicity mediated by immunologic synapse formation. Whether these molecules mediate cytotoxicity when secreted into the surrounding environment is unclear. In addition to soluble cytokines, classic innate-like effector molecules such as NKG2D could also potentially contribute to bystander killing (33). Indeed, initial experiments suggested that NKG2D could be induced on FAP-CAR-T cells following antigen engagement (data not shown). It is likely that bystander killing by CAR-T cells is complicated and multifactorial, with the contribution of different effector mechanisms varying according to such factors as the tumor type, the CAR-T cell phenotype and the susceptibility of individual target cells. Further research is warranted to better understand these mechanisms and how they are regulated. Indeed, if bystander killing can be optimized, uniform target antigen expression may not be required for efficient and complete tumor destruction by CAR-T cells. The potential of this approach has recently been observed using inhibitor of apoptosis protein (IAP) antagonists to facilitate antigen nonspecific killing by CAR-T cells targeting EGFRvIII ^47^.

The fact that bystander killing does not rely on antigen-directed targeting raises questions of safety, because there is a theoretical risk of targeting healthy cells and tissues. However, no overt toxicity was induced by FAP-CAR-T cells in our mouse xenograft model, even though the MO38 scFV has been shown to cross-react with murine FAP^24^. Moreover, co-culture experiments demonstrated that FAP-CAR-T cells can induce specific targeting of FAP^+^ tumor cells and bystander killing of FAP^-^ tumor cells while sparing non-malignant ECFCs within the same culture well. Together, these observations suggest that the bystander killing effector mechanisms operative in FAP-CAR-T cells may only be effective against cancer cells that express cognate receptors, whereas healthy cells lacking receptor expression are not targeted.

A typical CAR-T cell product contains both virally transduced (CAR-expressing) T cells and a variable proportion of non-transduced T cells. Here, we postulate that the non-transduced fraction of a CAR-T cell product can significantly contribute to killing of both antigen-positive and antigen-negative tumor cells through the process of bystander activation’. This involves activation of T cells in a T-cell receptor-independent and cytokine-dependent manner, and has long been observed during viral and bacterial infection^48, 49^, as well as in anti-tumor immunity ^29^. IL-18 and IL-15 are important factors that induce bystander activation of T cells ^50^, and IL-2 and IL-15 can convert T cells to lymphokine-activated killer (LAK) cells which exert cytotoxicity through antigen-independent mechanisms^31, 32^.

Of particular interest is IL-2, which was the first cytokine approved for human cancer therapy ^51^. IL-2 is a master cytokine which shapes the proliferation, differentiation and function of T cells and which can regulate the cytolytic machinery of T cells, including upregulation of perforin, granzymes and other proteins required for cytolytic granule fusion ^52^. In this study, we hypothesize that when stimulated by antigen, CAR-T cells can produce cytokines such as IL-2, which will induce the non-transduced T cells to exert cytotoxicity via antigen-independent mechanisms. Our observation of enhanced *in vitro* cytotoxicity achieved by supplementing with additional IL-2 may not appear novel on its own. However, this specific experiment offers us the opportunity to analyze various cytokines obtained from the assay and establish a correlation between cytokines and cytotoxicity. From this analysis, we have been able to determine which cytokines are likely to contribute to bystander killing and which are unlikely to do so (Figure 5c and Table 1). This IL-2-dependent function has the potential to broaden the antitumor effect of CAR-T products.

To mimic the heterogeneity of glioblastoma *in vivo*, we developed a mixed tumor model in which fluorescently tagged U87 and U251 cells were combined in an optimized ratio to produce tumors that had both cell types still present at endpoint. Using this model, we have shown that FAP-CAR-T cells can significantly delay tumor growth and extend survival, despite antigen heterogeneity. Complete tumor destruction was not observed in these experimental conditions, but it is worth noting that we used a low dose of CAR-T cells (1 x 10^6^) with relatively low transduction efficiency (∼25%) administered in a single intravenous dose. In future studies, a higher CAR-T cell dose with greater transduction efficiency may yield more complete tumor control. Analysis of tumor tissues at endpoint revealed that both U87 FAP^+^ and U251 FAP^-^ cells were reduced following FAP-CAR-T cell treatment, suggesting that the CAR-T cells exert longitudinal control over both U251 and U87 cells, in keeping with our *in vitro* findings of bystander killing. In these tumors, there were large areas of non-fluorescent tissue, constituting 90% of the area in the FAP-CAR-T treated group, 65% in the NT-T group, and 50% in the untreated group. These non-fluorescent cells may come from two sources: nonfluorescent tumor cells that finally escape immune control or infiltrating mouse-derived stromal cells such as fibroblasts and macrophages ^53^.

Our observations suggest that bystander killing mechanisms may enable CAR-T cells to target more tumor cells than would be predicted from the target antigen expression pattern, and thus may contribute to tumor destruction even in the presence of tumor heterogeneity. Approaches to boost endogenous bystander killing pathways may therefore increase the potency of CAR-T cell therapies, although the benefits of such approaches must be balanced against the theoretical risk of off-tumor toxicity. However, receptors such as NKG2D and even secreted cytokines are expected to act in a highly localized fashion. It was estimated, for example, that under physiologic conditions in dense tissues such as lymph nodes, the cytokine IL-2, interacts over a characteristic length scale of 8-14 cell diameters (80-140 µm), which is determined by a balance between diffusion and consumption of cytokine ^54^. Therefore, we might expect that bystander killing effects, may be limited to the tumor microenvironment where CAR-T cells engage with antigen.

In this study, we have developed a novel FAP-targeting CAR and shown, for the first time, that FAP-CAR-T cells effectively destroy glioblastoma cells *in vitro* and *in vivo*. These findings help to address the critical need for new and better target antigens for CAR-T cell therapy in the setting of glioblastoma. In addition, we highlight two mechanisms that broaden the antitumor effect of our CAR-T cell product: bystander killing of antigen-negative tumor cells, and cytokine-mediated bystander T-cell activation. Future work to better understand these mechanisms may reveal opportunities to enhance CAR-T cell function and promote more effective clearance of heterogenous solid tumors, a strategy that may be further augmented by multi-antigen targeting.

## METHODS

### Tumor cell lines

Long-term cell lines and GNS cells used in this study are listed in Supplementary Table 1. Short-term cultured GNS cell lines were a kind gift from Professor Bryan Day (QIMR Berghofer Medical Research Institute, Brisbane, Australia) or were generated in our laboratory as described^21, 55^. U251, U87 and RPMI-7951 cell lines were obtained from ATCC. U251-GFP and U87-RFP cells were generated via lentiviral transduction of parental cell lines with the lentiviral vectors pBLIV-MSCV-GFP-T2A-Fluc and pBLIV-CMV-RFP-T2A-Fluc (System Biosciences Inc, Palo Alto, USA), respectively. GFP- or RFP-expressing cells were sorted using MoFlo Astrios Cell Sorter (Beckman Coulter, Brea, USA). U251 and U87 (and their GFP/RFP-labelled variants) were maintained in Minimum Essential Medium Eagle supplemented with 10% fetal bovine serum (FBS), 1% penicillin/streptomycin, 1% glutamax, 1% sodium pyruvate, and 1% MEM non-essential amino acid solution (all Thermo Fisher Scientific, Scoresby, Australia). RPMI-7951 cells were maintained in RPMI-1640 medium supplemented with 10% fetal bovine serum (FBS), 1% penicillin/streptomycin and 1% glutamax. GNS cell lines were cultured in StemPro Neural Stem Cell medium (StemPro NSC, Thermo Fisher Scientific) in flasks, which were pre-coated with Matrigel (Corning, Silverwater, Australia) diluted 1/100 in PBS according to the manufacturer’s recommendations. Cultures were passaged using Accutase (Thermo Fisher) when they reached ∼70-90% confluence.

### CAR transgene design and generation of CAR-T cells

Anti-FAP scFv with Myc tag was obtained by PCR using donated plasmid containing MO36 as a template. The primers were F: GCCCGACGTCGCCACCATGGACTGGATCTGGCGCATC; R: TGCGGCCCCATTCAGATCCTCTTCTGAGATGAGTTTTTGTTCTGCGGCCGCCCGTTTTATTTCCAGC. A human IgG4 hinge/CH2CH3 was chosen as a long spacer to reduce the affinity to human FcRs^56, 57^. Aiming to avoid activation-induced cell death by further reducing its affinity to human and murine FcRs, additional modifications were made to the CH2 region. Three amino acid (PVA) derived from IgG2 substituted for four amino acids (EFLG) in the corresponding region of IgG4, and another glycosylation motif was mutated from N to Q^58, 59^. The CD3ζ and CD28 signaling domains were positioned after the spacer. Hinge, spacer and signaling domains were synthesized by GeneArt (Thermo Fisher). All fragments were assembled on pEntry vector and subcloned into lenti vector pPHLV-A.

The lentivirus encoding FAP-CAR was generated in HEK293T cells through transfection. HEK293T cells in a T175 flask were transfected with 46.4µg DNA consisting of equal amounts of four plasmids using 138µL lipofectamine 2000 (Thermo Fisher). The plasmids were: (i) pHLVA-FAP-CAR, which provides the lentiviral RNA containing the CAR (described in previous section); (ii) pMD2.G, which provides expression of VSV-G; (iii) pMDLgpRRE, which provides expression of gag pol; and (iv) pRSV-REV which provides the rev gene for lentiviral packaging. The supernatant was harvested 48h after transfection, centrifuged at 500 x *g* for 10min. and filtered through a 0.45µm filter (Nalgene Rapid-Flow™, Thermo Fisher), ready for transduction. To produce CAR-T cells, a 24-well plate was pre-coated with anti-CD3 (OKT3) and anti-CD28 antibodies (15E8) (Miltenyi Biotec, Macquarie Park, Australia) at 1µg mL^-1^ on day 0. PBMC were seeded at 1 × 10^6^ cells/well on day 1 in Advanced RPMI (Thermo Fisher) with 10% FBS, with IL-7 and IL-15 (Miltenyi) added at 10 ng mL^-1^ and 5 ng mL^-1^, respectively, on day 2. On day 3, the activated T cells were transduced by spinoculation with lentivirus encoding FAP-CAR. In brief, 0.5mL T cells at 2 × 10^6^ cells mL^-1^ were mixed with 1.5mL lentiviral supernatant and centrifuged in 24-well plates pre-coated with 7 µg mL^-1^ Retronectin (Takara Bio, San Jose, USA) at 1000 x *g* for 20min. The transduced T cells were expanded for another 4 days with IL-7/IL-15 before cryopreserving in FBS + 10% DMSO and storage in liquid nitrogen. Before use, the CAR-T cells were thawed and rested overnight in Advanced RPMI with 10% FBS. CAR expression was detected by FITC-conjugated anti-Myc tag antibody (Abcam, Melbourne, Australia) and analyzed by flow cytometry using an Accuri C6 Plus flow cytometer (BD Biosciences, Macquarie Park, Australia).

### Flow cytometry analysis of FAP expression

U251, U87 and RPMI-7951 cells were stained by PE-conjugated anti-FAP antibody (clone 427819, R&D Systems) in 5% BSA/PBS staining buffer for 30min. Cells were washed with staining buffer and 1µL 7AAD (Thermo Fisher) was added in a final resuspension volume of 200µL before analysis by flow cytometry (BD Accuri™). For staining of GNS cultures, cells were harvested using Accutase (Thermo Fisher), washed in PBS and resuspended in FACS buffer (PBS + 1% bovine serum albumin + 0.04% sodium azide). Cells were then stained as described above.

**T-Cell Phenotype:**

T cells were resuspended in 200μL of FACs buffer at a concentration of 2 × 10^6^ cells mL^-1^. Human Fc block (BD) was added (2μL per sample) and incubated for 5min on ice, then samples were centrifuged at 300 × g for 5min and the pellets resuspended in 50μL of FACs buffer. Samples were stained for 30min on ice using the following antibodies: anti-CD3-PE (BD), anti-CD4 BV510 (BD), anti-CD8 PE-Cy7 (BD), anti-CD45RA APC (BD), anti-CCR7 (CD197) BV421 (BD), anti-Myc AF488 (Abcam). Samples were washed twice with FACs buffer before analysis by flow cytometry using the Accuri C6 Plus, as per above.

### Cytotoxicity assays

Four different cytotoxicity assays were used to address different biologic questions.

#### Cytotoxicity assay using real-time impedance-based cell analysis

The Axion BioSystems (Atlanta, USA) Maestro Z system was used to measure cytotoxicity. After addition of 100µL of culture medium to the CytoView-Z plate (Axion Biosystems), a baseline reading was taken and then target cells were seeded in a volume of 100µL at 15,000 cells/well for U87 and U251 or 10,000 cells/well for GNS cultures. The wells were pre-coated with Matrigel if GNS cells were used as targets. Resistance values were monitored every 60sec using AxisZ software during overnight culture at 37℃ in the Maestro Z instrument. Once formation of a stable monolayer was confirmed, 100µL of CAR-T cells were added in the same medium as the target cells. Resistance values were normalized to a time shortly after adding CAR-T cells once readings had re-stabilized. Cytotoxicity was measured as %lysis based on the impedance ratio of tested cells and untreated control cells, which was calculated by AxisZ software.

#### Cytotoxicity assay by flow cytometry

GNS cells (1 x 10^5^) were co-cultured with an equal number of FAP-CAR-T cells or control non-transduced T cells (NT-T) in equivalent culture conditions in StemPro NSC medium in a 24-well plate. In co-cultures of pairs of GNS cell lines of differing levels of FAP expression, FAP-expressing GNS cells were first pre-labeled with CellTrace™ Far Red (CTFR; Thermo Fisher) at 1µM for 20min at 37^0^C, to differentiate the populations, and equal numbers of FAP^+^ and FAP^-^ cells were mixed. After 48 hours, T cell suspension were collected and adhered tumour cells were detached and harvested using StemPro™ Accutase™ (Thermo Fisher). T cells and tumor cells were pooled together and collected by centrifugation. Cells were stained in PBS with Fixable Viability Stain 575V (BD) or Fixable Viability Dye eFluor™ 506 (Invitrogen) where indicated for 10min at room temperature, before antibody staining. The antibodies used for staining included: BB515 anti-Fas (clone DX2, BD Biosciences), BV786 anti-CD3 (clone SK7, BD Biosciences), PE anti-FAP (clone 427819, R&D Systems) and BV480 anti-FasL (clone NOK-1, BD Biosciences). Samples were acquired on an LSR Fortessa (BD Biosciences) and data analyzed using FCS Express 7 Cytometry (De Novo software, Pasadena, CA, USA).

#### Cytotoxicity assay to assess the role of soluble factors

These experiments were conducted in 24-well Transwell plates (Corning) or Nunc Cell Culture Inserts (Thermo Fisher) with a 0.4µm pore size. FAP-expressing cells were plated in the lower well at 1 x 10^5^ cells/well, and 1 x 10^4^ FAP-negative cells were seeded in the upper inserts. When monolayers of target cells had established, FAP-CAR-T cells or non-transduced T cells were added to the bottom well at a 1:1 E:T ratio. After 48h incubation at 37°C, cells in the inserts were stained using propidium iodide (PI) (Thermo Fisher) at 1 µg mL^-1^ and Hoechst 33342 (Thermo Fisher) at 2µM for 20min at 37°C. Without washing, inserts were imaged using an IX73 microscope equipped with CoolLED pE-4000 light source and running CellSens software (Olympus). The cells on each image were counted by QuPath (University of Edinburgh). Cell viability was calculated as [1 - (PI counts/Hoechst counts)] x 100%.

#### Cytotoxicity assay using fluorescence imaging of U87-RFP and U251-GFP cells

U87-RFP and U251-GFP were seeded with starting numbers of 5× 10^4^ cells/well. FAP-CAR-T cells were immediately added at a 1:1 E:T ratio. The cocultured cells were imaged by the Olympus IX73 microscope 24h later using 100ms exposure for RFP and 3ms exposure for GFP. Three fields of views were taken from each well and duplicate wells were used for one assay. The images were analyzed by ImageJ for fluorescence counts using threshold values of 68-255 for RFP and 98-255 for GFP. All images were collected and analyzed using the same parameters for consistency.

#### Endothelial Colony Forming Cell (ECFC) co-culture experiments

Endothelial colony-forming cells (ECFCs) were derived from peripheral blood obtained from healthy human donors, following established protocols^60, 61^. Briefly, peripheral blood mononuclear cells were plated on collagen I (Corning) -coated 48 well plates in EGM-2 medium (Lonza, Switzerland), supplemented with 20% ES cell-screened FBS (Hyclone, GE Healthcare, Chicago, USA) until colonies formed. Cells were passaged and maintained for no more than nine passages in EGM-2 medium with 20% FBS, with confirmation of ECFC markers using flow cytometry. A mixed target cell population consisting of ECFCs, U87-RFP, and U251-GFP cells (1.5 X 10^5^ each) was co-cultured for 48 hours in EGM-2 medium with 6 X 10^5^ effector cells in 24-well plates. Subsequently, the cells were washed, treated with trypsin, harvested in 1.5 mL PBS, and analyzed using an Accuri C6 Plus flow cytometer. From each sample, cells from a fixed volume of 66 µL were counted. U87-RFP and U251-GFP cells were counted based on fluorescent gating, while ECFCs were identified through their lack of fluorescence. The cell counts obtained after 48 hours were compared with those from samples collected at 0 hours to determine the fold change.

### Measurement of secreted factors

To measure IL-2 secretion, target cells were seeded in a flat-bottom 96-well plate at 20,000 cells/well on day 0. On day 1, 1×10^5^ CAR-T cells were added per well. The supernatant was collected on day 3 and analyzed by human IL-2 ELISA Kit (Thermo Fisher) according to the manufacturer’s protocol.

To measure levels of 13 soluble factors simultaneously using the LEGENDplex Human CD8/NK Panel (BioLegend, San Diego, USA), supernatants were collected from cytotoxicity assays conducted in the Maestro Z instrument at endpoint and stored at -20°C. Assays were conducted according to the manufacturer’s protocol and analyzed using a BD LSR Fortessa.

### Murine subcutaneous tumor xenograft model

Animal experiments were approved by the University of South Australia Animal Ethics Committee (Ethics number: 45/17). Female nonobese diabetic/severe combined immunodeficiency/IL2Rγ-null (NSG) mice were purchased at 6 weeks from the Animal Resource Centre (Perth, Australia) and housed in the Immunocompromised Suite of the Core Animal Facility of the University of South Australia in isolator cages. Mice were provided with standard chow and sterile water *at libitum*, and given soaked food and sunflower seeds as required to support weight maintenance. At least 2 weeks were allowed for acclimatization prior to initiating experiments. Sample size for treatment and control groups was calculated *a priori* based on f = 0.40, SD = 0.2, power level 0.9 for the primary outcome measure of tumor volume. Mice were randomly allocated to groupings (n = 4-5 per cage) at the beginning of the experiment; no blinding was performed. To minimize potential confounders, animals were housed in adjacent cages under the same conditions (lighting, food, handling, etc). To create tumors of mixed cell populations, an admixture of 1 × 10^6^ U251-GFP cells and 2 × 10^6^ U87-RFP cells were injected subcutaneously in both flanks. Tumors were allowed to establish for 11 days and then 1 × 10^6^ CAR-T cells were administered by tail vein injection. If one or both tumors failed to establish, that animal was excluded from the experiment (n = 1). Mice were monitored daily until tumors became palpable, after which time tumor growth was measured using an electronic caliper every 2-3 days. The volume of the tumor was calculated as length × width^2^ ×1/2. Mice were humanely killed when at least one tumor volume reached 1000mm^3^. Tumors were harvested at the experimental end point and prepared for frozen sections. Tumor samples fixed in 10% buffered formalin overnight were cryoprotected with 10% sucrose to 30% sucrose in PBS sequentially before embedding in OCT (Sakura Finetek, Tokyo, Japan) and freezing on dry ice, with subsequent storage at -80°C. This protocol was selected and optimized to allow maximum retention of soluble fluorescent proteins (GFP/RFP) during tissue processing. Cryosections (5µM) were cut using a Leica CM1950 cryostat and mounted using ProLong Gold with DAPI (Thermo Fisher). Slides were imaged using the Olympus IX73 microscope. To minimize bias, fields of view were first selected based on DAPI nuclear staining, before imaging the corresponding GFP and RFP channels. Areas of green and red fluorescence were automatically calculated using ImageJ (NIH). Threshold 21-255 was used for RFP area calculation while threshold 35-255 was used for GFP area calculation. Fluorescence overlays were created by overlaying black and white images and applying false colors using ImageJ.

### Statistical analysis

Statistical analyses were performed using GraphPad Prism version 5 and OriginPro 8 (OriginLab). In addition, the TumGrowth web program (https://kroemerlab.shinyapps.io/TumGrowth/) was used for comparison of tumor growth curves by Type II ANOVA. For Kaplan-Meier Survival analysis, Log Rank method was used to test the Equality over Groups.

## Acknowledgments

We thank Professors Klaus Pfizenmaier (Institute of Cell Biology and Immunology, University of Stuttgart, Stuttgart, Germany), and Stephen Gottschalk (Department of Bone Marrow Transplantation and Cellular Therapy, St Jude Children’s Research Hospital, Memphis, TN, USA), for provision of the FAP-specific MO36 scFv. We are grateful to Professor Bryan Day (QIMR Berghofer Medical Research Institute, Brisbane, Queensland, Australia) for the IIIA4 single chain variable fragment used for construction of the EphA3-specific CAR. We also thank Dr Susanne Heinzel, Walter and Eliza Hall Institute of Medical Research, Melbourne Victoria, Australia, for her review and valuable comments on the manuscript.

## DECLARATIONS

### Ethics approval and consent to participate

Blood collection from healthy donors for CAR-T cell manufacture was approved by the Central Adelaide Local Health Network (CALHN) Human Research Ethics Committee (approval number HREC/16/RAH/191). Blood collection from healthy donors for ECFC generation was approved by the human ethics committees of SA Pathology and the University of South Australia (approval numbers, ethics #85–13 and UniSA HREC #201187 respectively). Written consent to participate was obtained from donors of fresh whole blood, whereas buffy coats obtained from Australian Red Cross Lifeblood were obtained under an approved waiver of consent. Animal experiments were approved by the University of South Australia Animal Ethics Committee (approval number: 45/17).

### Consent for publication

All authors consent to the publication of this manuscript.

### Availability of data and materials

Original datasets will be made available upon reasonable request to the corresponding author.

### Authors’ contributions

**WY:** investigation, methodology, conceptualization, data curation, formal analysis, resources, writing – original draft, writing – review & editing. **NTHT:** Investigation, methodology, formal analysis. **RP:** Investigation, formal analysis. **TG:** Investigation, formal analysis, supervision, writing – review & editing. **MNT:** Resources. **SMP:** Funding acquisition, resources. **MPC:** Resources. **CSB:** Resources. **LME:** Conceptualization, investigation, methodology, data curation, formal analysis, funding acquisition, resources, supervision, writing – original draft, writing – review & editing. **MPB:** Conceptualization, funding acquisition, resources, supervision, writing – review & editing.

