## Supplementary table and figures for "Endogenous bystander killing mechanisms enhance the activity of novel FAP-specific CAR-T cells against glioblastoma"

### Supplementary Table 1

| Cell Name | Type | Antigen | Notes |
|---|---|---|---|
| RPMI 7951 | Melanoma cell line | $FAP^{+}$ high | For FAP-CAR-T quality control |
| U251 | Glioblastoma cell line | $FAP^{-}$ | |
| U87 | Glioblastoma cell line | $FAP^{+}$ | |
| U251-GFP | Glioblastoma cell line | $FAP^{-}$ | For mouse model |
| U87-RFP | Glioblastoma cell line | $FAP^{+}$ | For mouse model |
| CCB-G3-C | GNS | $FAP^{+}$ high | |
| MN1 | GNS | $FAP^{+}$ medium | |
| CCB-G6-T | GNS | $FAP^{+}$ low | |
| RKI-1 | GNS | $FAP^{-}$ | |
| SJH-1 | GNS | $FAP^{-}$ | |
| CCB-G25-T | GNS | $GD2^{+}$ | Test bystander killing on alternative target |
| HW1 | GNS | $GD2^{+}$ | Test bystander killing on alternative target |
| HEK293t | Human embryonic kidney | | For lentivirus production |

Supplementary Table 1: Cell lines and GNS cells used in this study

### Supplementary Figure 1

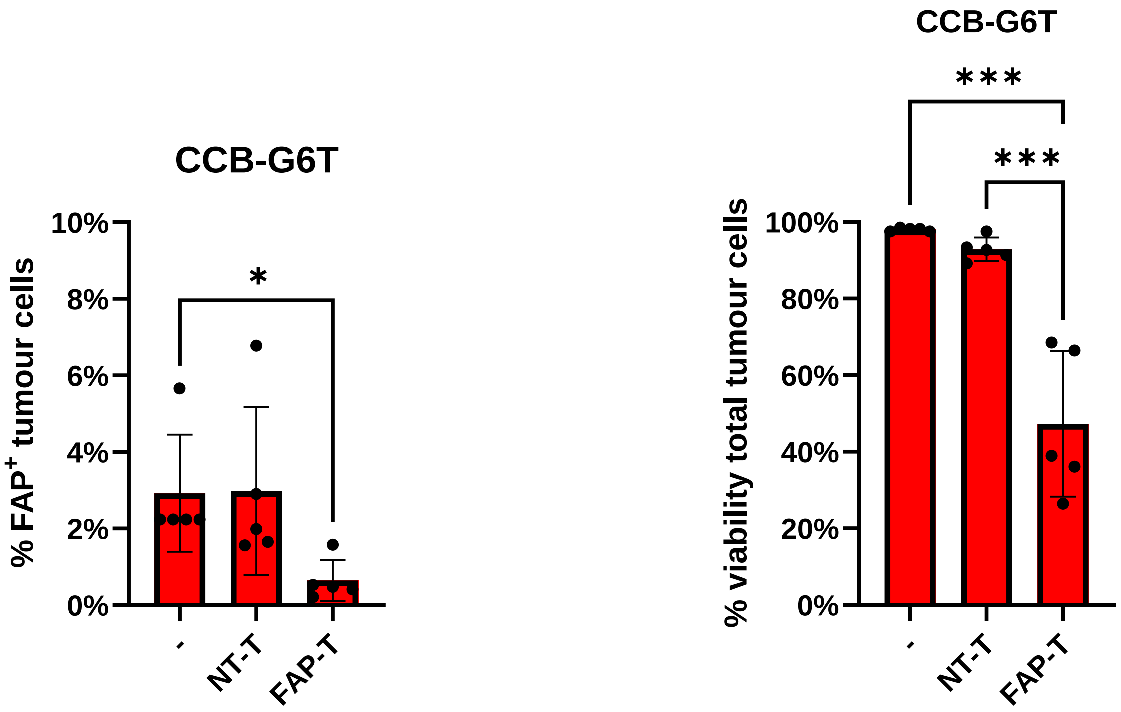


**Supplementary Figure 1. CCB-G6-T and CAR-T co-culture.** FAP expression (left) and viability (right) of the CCB-G6-T GNS cell line was analyzed after 48 hours co-culture with FAP-CAR-T cells or NT-T cells. Data are pooled from 5 experiments using 3 different PBMC donors for CAR-T cell manufacture. Paired *t*-test * $P < 0.05$, *** $P < 0.001$.

### Supplementary Figure 2

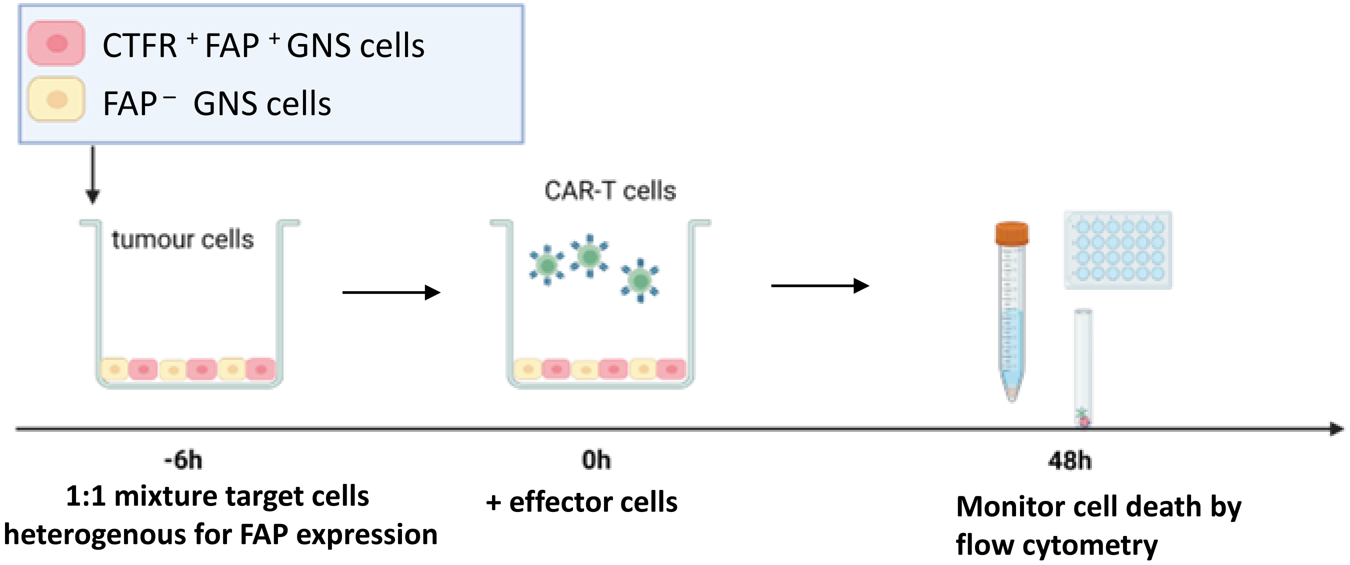


**Supplementary Figure 2: Schema illustrates the workflow of co-culture assay**: $FAP^{+}$ and $FAP^{-}$ tumor cells were mixed at 1:1 ratio and treated with either FAP-CAR-T cells or non transduced T cells (NT-T) at 1:1 E:T ratio. As a control, single cultured target cells were treated in the same way (not illustrated, Figure S3). The viability of target cells was analyzed 48h later by flow cytometry.

### Supplementary Figure 3

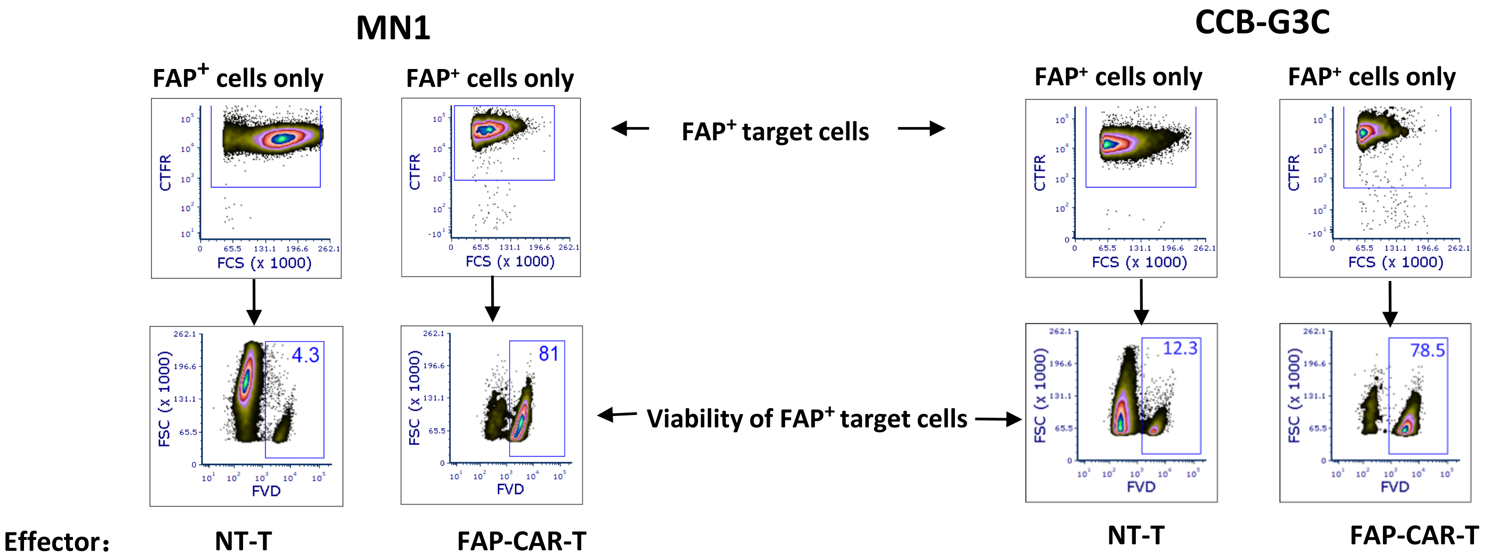


**Supplementary Figure 3: Representative flow cytometry data (from experiments shown in Figure 3), demonstrating that FAP-CAR-T cells efficiently kill FAP+ tumor cells:** Representative analysis of MN1 (Pair 1) and CCB-G3C (Pair 2) is shown here. Similar results were observed in all experiments.

### Supplementary Figure 4

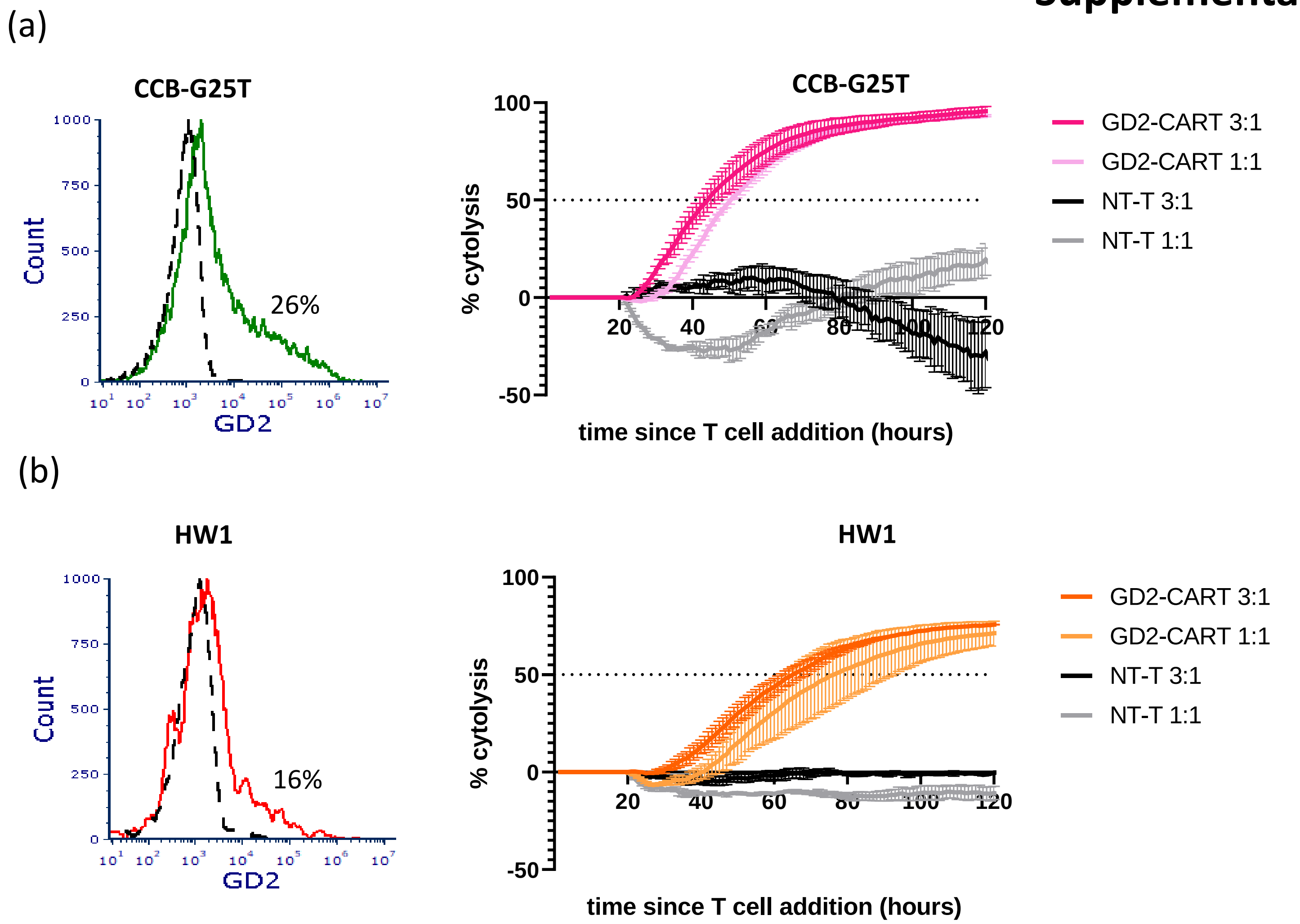


**Supplementary Figure 4: Real time cytotoxicity assays of GD2 CAR-T cells demonstrate potential bystander killing function.** Low-level GD2-expressing GNS cell lines (**a**: CCB-G25T; **b**: HW1) were cultured in a CytoView-Z plate for 48h before addition of GD2 CAR-T cells or NT-T cells as control. The E:T ratios are indicated. Target cell growth was monitored by impedance values in real time by Maestro Z system. The percent of target cell cytolysis was determined by the ratio of impedance values of treated and untreated cells at the same time. Duplicate wells were averaged for each experimental condition. The curve represents the mean while the bar represents the SEM.

### Supplementary Figure 5

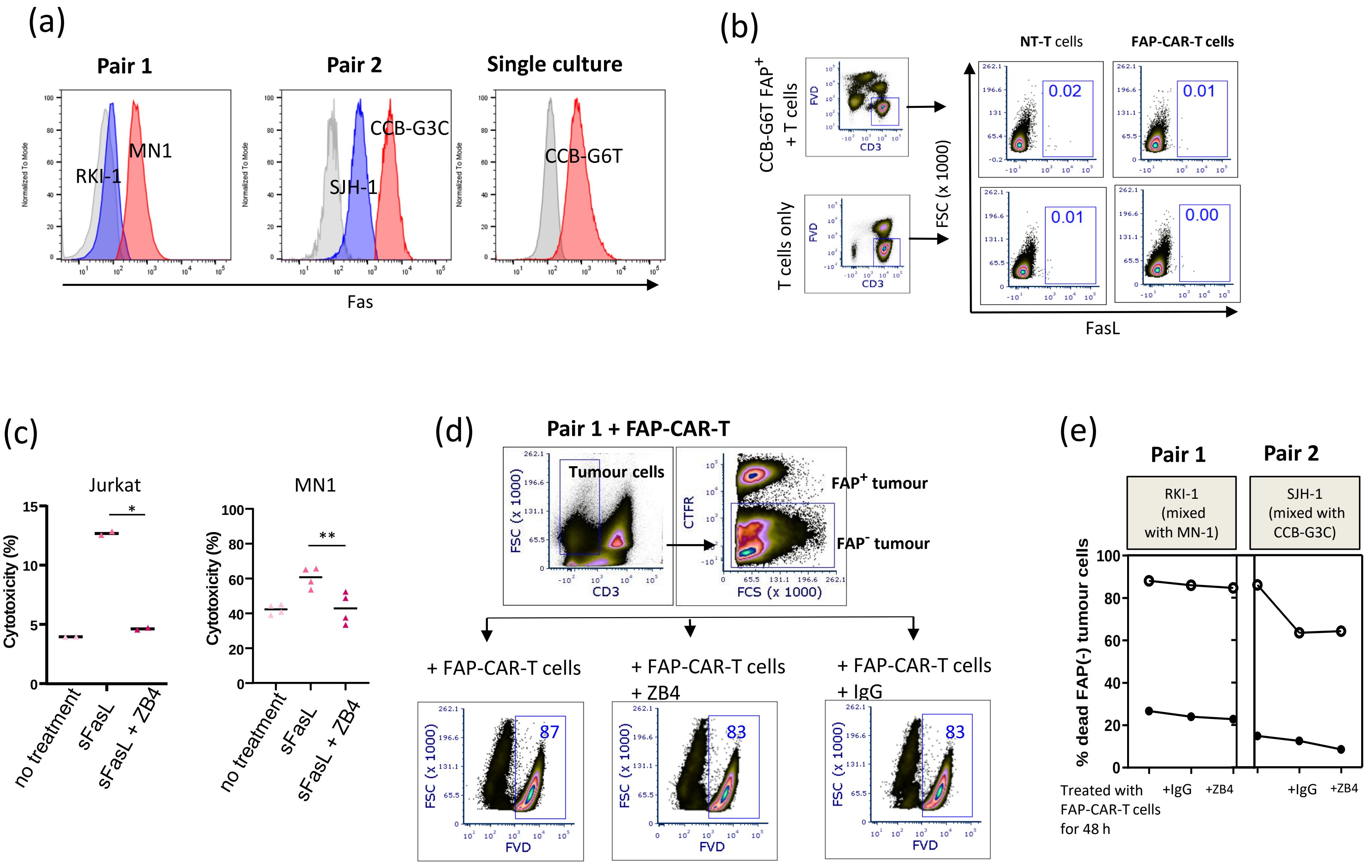

**Supplementary Figure 5: Fas/FasL expression and Fas blocking: (a)** mean fluorescence intensity (MFI) of surface Fas expression on GNS cell lines used as target cells in Fas blocking assays. Histograms represent two pairs of mixtures: RKI-1 $FAP^-$ and MN-1 $FAP^+$, or SJH-1 $FAP^-$ and CCB-G3C $FAP^+$, or CCB-G6-T (partial $FAP^+$). Grey histogram represents the isotype control of Fas staining. Data are representative of 3 experiments. **(b)** Representative FasL expression of NT-T cells or FAP-CAR-T cells before (lower panel) and after (upper panel) co-culture with CCB-G6T. Data are representative of 3 experiments. (**c**) Jurkat cells (n = 2) or MN-1 cells (n = 4) were incubated with ZB4 antibody to block FasL (2.5 µg $ml^{-1}$) or left untreated for 1 hour before being supplemented with sFasL (200 ng $ml^{-1}$). Cells were collected after 24 hours, stained with Hoechst (15 µM) and PI (1 µg $mL^{-1}$) for 45 minutes at RT and then analyzed by flow cytometry. MN-1 cells were pre-sensitized with 4mM sodium butyrate (present in culture). *t*-test, * $P < 0.05$, ** $P < 0.01$ **(d)** Representative viability staining of RKI-1 cells in co-culture with MN-1 and FAP-CAR-T cells, in the presence of Fas blocking antibody ZB4 (middle plot) or control IgG (right plot). T cells were excluded by $CD3^+$ gating. **(e)** Viability change of $FAP^-$ GNS after 48h co-culture assay with ZB4 blocking. Isotype IgG was used as control. Filled and open symbols represent two independent assays carried out for Pair 1 group and Pair 2 group.

### Supplementary Figure 6

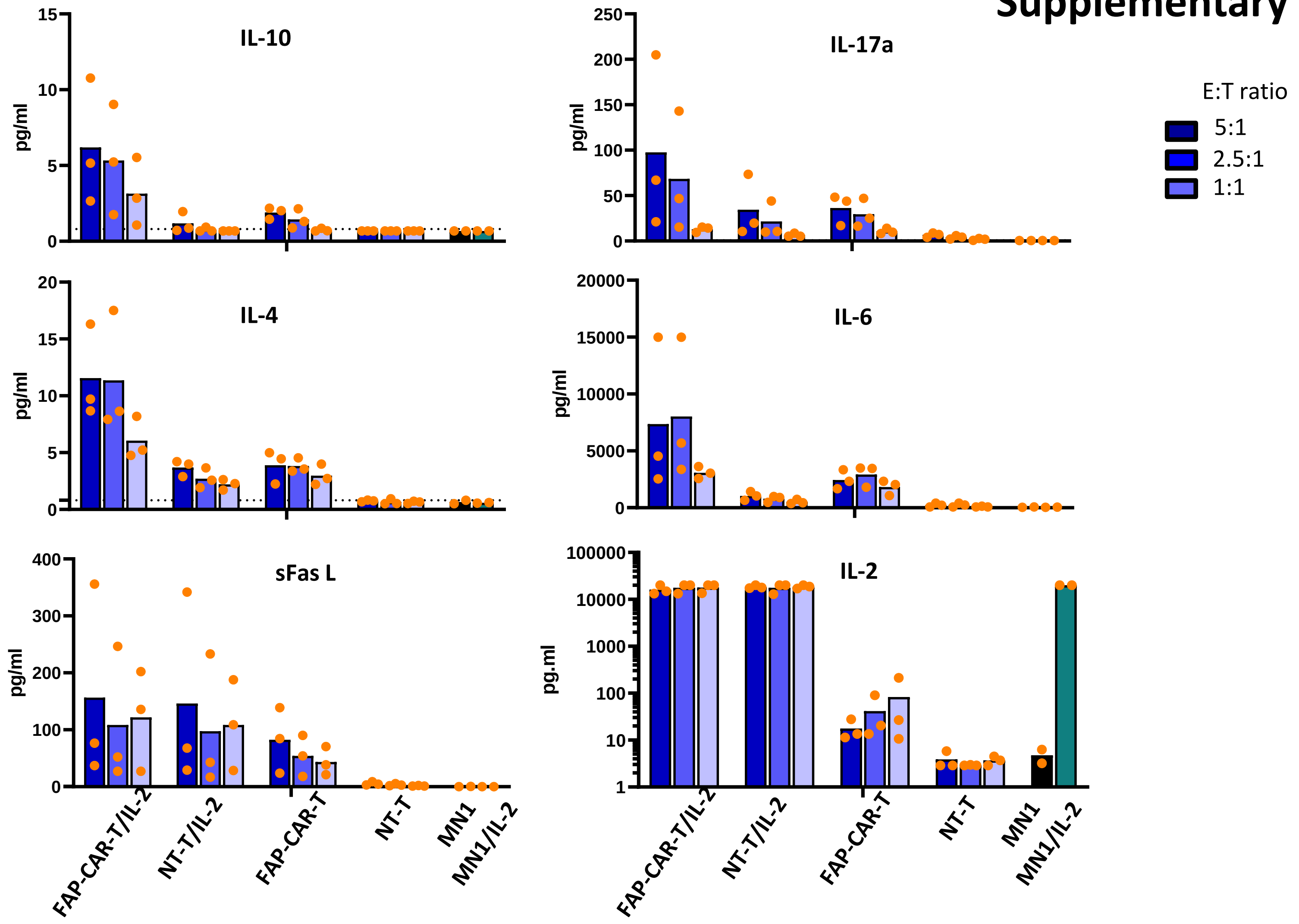


**Supplementary Figure 6: Cytokines produced during co-culture of FAP-CAR-T cells and GNS target cells, with and without addition of exogenous IL-2:** FAP$^+$ MN1 cells were established in a CytoView-Z plate for 24h prior to addition of effector cells. Target cell growth was monitored by impedance values in real time by the Maestro Z system and supernatants were collected for measurement of secreted factors 50h after adding effector cells. Levels of cytokines and other soluble factors were assessed using Legendplex assay. Results of 6 analytes are shown here (the other 6 were shown in Figure 5). Data are pooled from 3 experiments (or 2 experiments for MN-1 and MN-1/IL-2 controls).

### Supplementary Figure 7

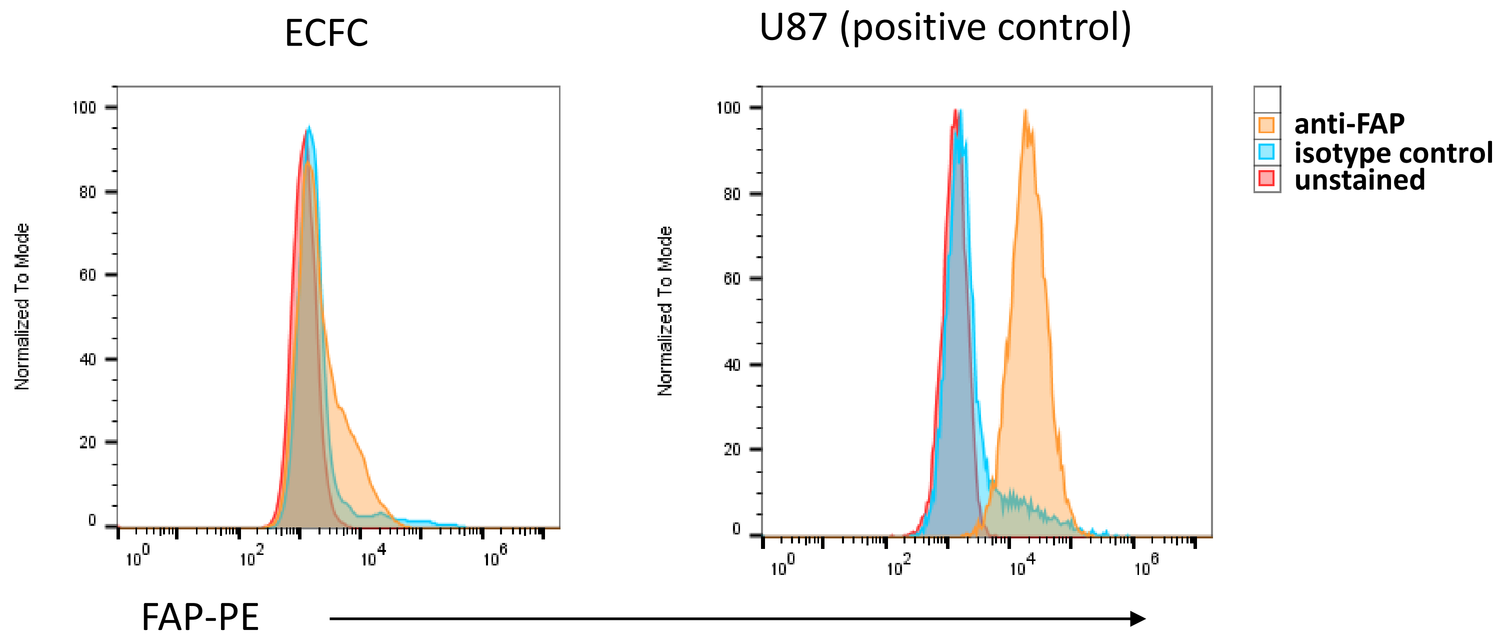


**Supplementary Figure 7: Healthy primary ECFC express minimal FAP:** ECFC were stained for flow cytometry using PE-conjugated anti-FAP antibody or isotype-matched control antibody, or left unstained. U87 cells were stained in the same manner on the same day, as positive control.

### Supplementary Figure 8

**(a)**

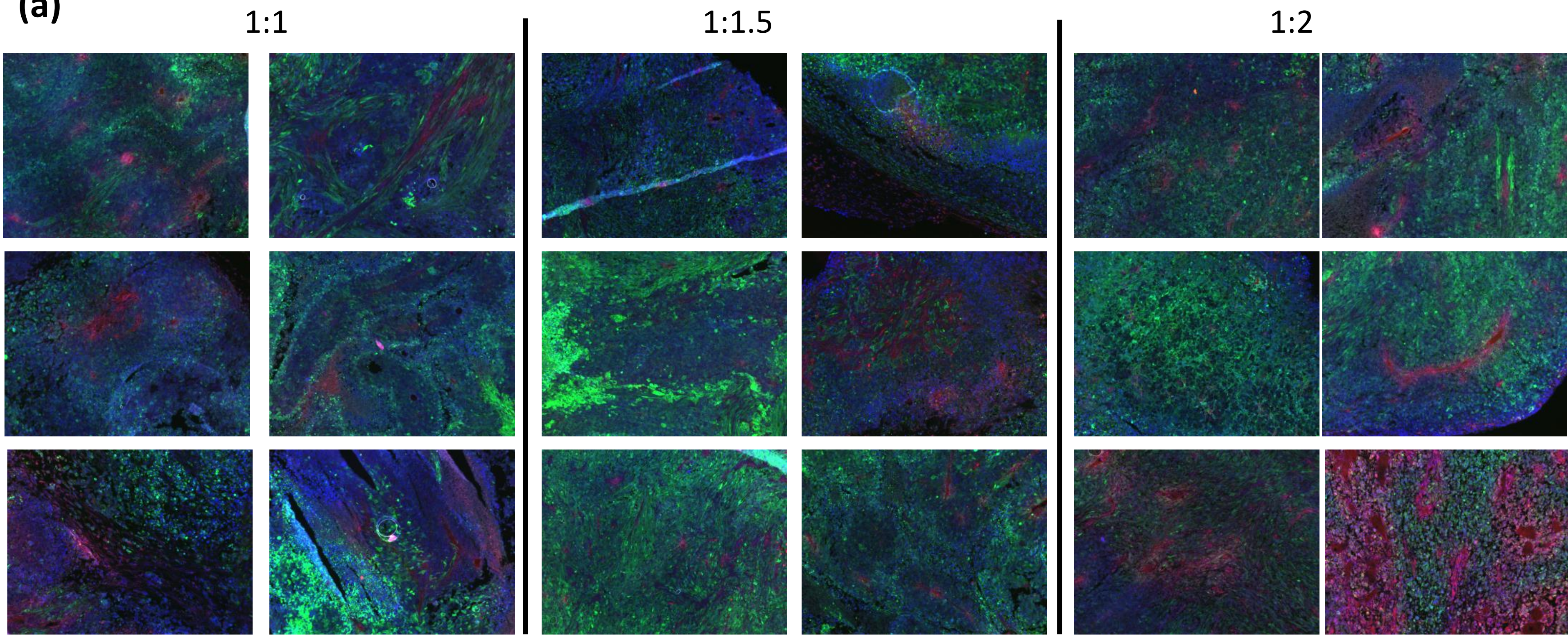


**(b)**

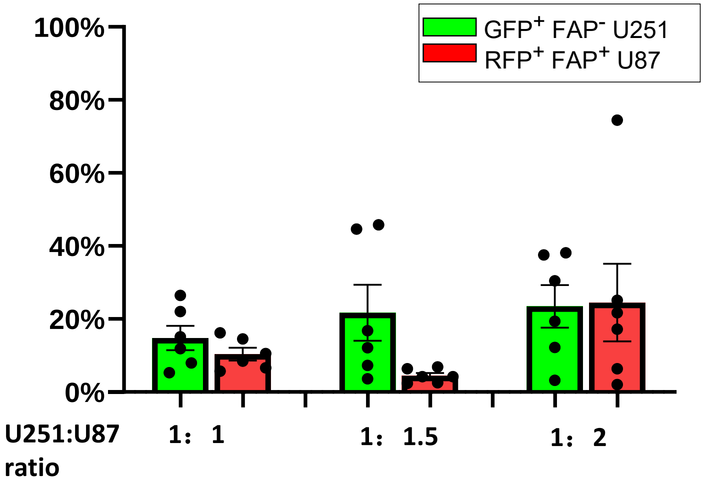


**Supplementary Figure 8. Optimizing the ratio of U251:U87 cells for the mixed tumour subcutaneous animal model. (a)** A mixture of $FAP^-$ U251-GFP cells and $FAP^+$ U87-RFP cells at 3 different ratios was injected subcutaneously on both flanks of NSG mice. Each injection contained $1\times10^6$ U251 cells with $1\times10^6$ (1:1), $1.5\times10^6$ (1:1.5), or $2\times10^6$ (1:2) U87 cells. The samples from each group were harvested when the tumor reached 1000 $cm^3$ and the cryosections were examined for GFP and RFP expression. **(b)** The % of area positive for GFP or RFP signal was determined using ImageJ.
